# Mixture of Learning Strategies Underlies Rodent Behavior in Dynamic Foraging

**DOI:** 10.1101/2022.03.14.484338

**Authors:** Nhat Minh Le, Murat Yildirim, Yizhi Wang, Hiroki Sugihara, Mehrdad Jazayeri, Mriganka Sur

## Abstract

In volatile foraging environments, animals need to adapt their learning in accordance with the uncertainty of the environment and knowledge of the hidden structure of the world. In these contexts, previous studies have distinguished between two types of strategies, model-free learning, where reward values are updated locally based on external feedback signals, and inference-based learning, where an internal model of the world is used to make optimal inferences about the current state of the environment. Distinguishing between these strategies during the dynamic foraging behavioral paradigm has been a challenging problem for studies of reward-guided decisions, due to the diversity in behavior of model-free and inference-based agents, as well as the complexities that arise when animals mix between these types of strategies. Here, we developed two solutions that jointly tackle these problems. First, we identified four key behavioral features that together benchmark the switching dynamics of agents in response to a change in reward contingency. We performed computational simulations to systematically measure these features for a large ensemble of model-free and inference-based agents, uncovering an organized structure of behavioral choices where observed behavior can be reliably classified into one of six distinct regimes in the two respective parameter spaces. Second, to address the challenge that arises when animals use multiple strategies within single sessions, we developed a novel state-space method, block Hidden Markov Model (blockHMM), to infer switches in discrete latent states that govern the choice sequences across blocks of trials. Our results revealed a remarkable degree of mixing between different strategies even in expert animals, such that model-free and inference-based learning modes often co-existed within single sessions. Together, these results invite a re-evaluation of the stationarity of behavior during dynamic foraging, provide a comprehensive set of tools to characterize the evolution of learning strategies, and form the basis of understanding neural circuits involved in different modes of behavior within this domain.

## Introduction

Reward-guided decision making has largely been studied in terms of two broad regimes of behavioral strategies and neural systems. One influential class of models involve reinforcement learning models in which each action has an internal value that is updated over time based on feedback from the environment^1,2^. Variants of these model-free approaches, such as the Rescorla-Wagner updating rule^3^, the Q-learning algorithm^4^, local matching strategies^5^, or Thomson sampling^6^, have been influential in formulating efficient decision-making and learning strategies in uncertain environments^7–12^. These models have also been successful in explaining the activity of cortical and subcortical areas in relation to reward prediction errors^13^, action values^7,14^ or previous choice and outcome history^15,16^.

When reward and outcome contingencies follow a specific structure and regularity, another set of models, inference-based models with trial-to-trial Bayesian updates, are often used to simulate the actions of agents^17–19^. This type of strategy involves the use of internal models to make efficient inferences about the hidden states and optimal actions. Such inference-based (also known as model-based) behavior are often seen only in expert animals that are familiar with the structure of the task and able to hold an internal representation and understanding of the dynamics of the surrounding world^17,18^. Inference-based behavior has also been shown to engage a non-overlapping set of brain areas from those that are involved in model-free strategies^20,21^.

In many previous studies of reward-guided decision-making, these two modes of behavior, model-free and inference-based learning, have largely been treated as separate behavioral domains that require different sets of analytical tools and models. For example, reinforcement learning models and logistic regression models have often been used in a subset of studies that assume a model-free structure of behavior^7,14^. This model-free approach allows researchers to answer questions related to the value representations in different brain areas, as well as study the effect of perturbations on the parameters of the models^15,22–24^. On the other hand, a complementary set of studies focus on the behavior of well-trained animal with the assumption that these animals behave exclusively in the inference-based domain^25,26^. While these stationarity assumptions are helpful when animal behavior exclusively belongs to one domain or another, recent studies have started to bring attention to the overlap and interaction between these types of strategies^19,27^. For example, it was found that in the same dynamic foraging task, rodents might engage in both model-free and inference-based behavior, transitioning from the former strategy to the latter with experience in the environment^17,18^. Another set of studies highlighted additional complexity in rodent behavior, as they often switch between states of engagement and disengagement during decision-making tasks^28,29^. These results suggest model-free and inference-based behavior might be interspersed within the same session, potentially engaging different subsets of neural circuits and mechanisms for parallel computation of multiple decision variables^30^. The use of mixture of strategies is further supported by the discovery of separable components of rodent behavior in a reward-guided task^27^. Together, these results call for a more unified approach for dissecting the two sets of strategies and understanding the transitions between them during learning as well as within single sessions of the task.

Here, we focused on the problem of distinguishing these two types of behavior in the dynamic foraging paradigm (also known as the two-armed bandit task), a standard behavioral framework of previous investigations into reward-guided behavior^31,32^. Our main goal is to develop a set of behavioral benchmarks, analytical tools and approaches to help reliably dissociate between the two classes of strategies. This is a challenging endeavor for two primary reasons. First, these two classes of models are qualitatively distinct in form: model-free approaches involve agents that update their action values from trial to trial with a learning rate and an exploration parameter^1^, while inference-based approaches involve agents with a prior and internal model specified by some parameters^33^. We are thus faced with two sets of parameters with which to fit the behavior, and will need to compare how well these parameter spaces can fit the same sequence of observations. The second analytical challenge occurs when animals mix between different modes of behavior in the same session. With this mixing, techniques that rely on aggregate measures of behavior over entire sessions will lead to inaccurate estimates of behavioral parameters, as we will show in our subsequent analyses, requiring alternative methods to segment and infer latent states of the behavior from trial to trial.

To present our approach for distinguishing between the two types of strategies in dynamic foraging, the paper is organized as follows. We first describe our experimental setup to study dynamic foraging behavior in head-fixed mice. To analyze the behavior of our animals during training, we focus on two models, (1) model-free agents that implement the *ε*-greedy Q-learning decision strategy, and (2) inference-based agents that hold a Markovian internal model of the world. With this formulation, we show that current analytical methods are inadequate to fully dissociate between the two classes of strategies, as these methods are insufficient to account for the diversity of learning across the parameter spaces. In addition, methods that rely on session-averaged metrics might give rise to inaccurate estimates of the behavior when animals mix between behavioral strategies. We then present our approach to overcome the two challenges. To comprehensively compare the behavior of the two models, we characterize four main behavior features of the agent’s switching dynamics and perform a complete survey of these features across the inference-based and Q-learning parameter spaces. This analysis reveals distinct behavioral clusters which can be robustly decoded from each other, with a decoding accuracy close to 100% between model-free and inference-based agents. To address the difficulty of behavioral analysis of mixtures of strategies, we have built a novel state-space model (blockHMM) to infer the latent states of behavior sessions, eliminating the potential confound of mixtures of learning strategies on behavioral analysis. We validate this approach with simulations to demonstrate its reliability in recovering the hidden states of behavior from observed choice sequences. Together, these new tools reveal the highly dynamic nature of rodent behavior in this task, further highlighting the variabilities between animals and the need for a statistical approach based on inferred latent states for understanding the structure of task behavior.

## Results

### Dynamic foraging task and decision strategies of model-free and inference-based agents

We trained head-fixed mice on a dynamic foraging (two-armed bandit) task (Fig. 1a). Mice were placed on a vertical rotating wheel^34^, and on each trial, they were trained to perform one of two actions, left or right wheel turns. On each trial, one movement was rewarded with probability of *p* and the other with the complementary probability of 1 – *p*. We tested mice in different dynamic environments with different values of *p*. In the ‘100-0’ environment, one action yielded reward with 100% probability, while the alternative yielded no reward (Fig. 1b). Similarly, in ‘90-10’, ‘80-20’ and ‘70-30’ environments, reward probabilities were assigned to the two indicated values. The environments were volatile such that the high- and low-value sides switched after a random number of trials sampled between 15-25 without any external cues, requiring agents to recognize block transitions using only the reward feedback. To ensure stable behavioral performance, we also required the average performance of the last 15 trials in each block to be at least 75% before a state transition occured. We collected behavioral data from *n* = 21 mice that were trained in the task for up to 40 sessions per animal (typical animal behavior shown in Fig. 1c for a 90-10 environment).

**Figure 1.**
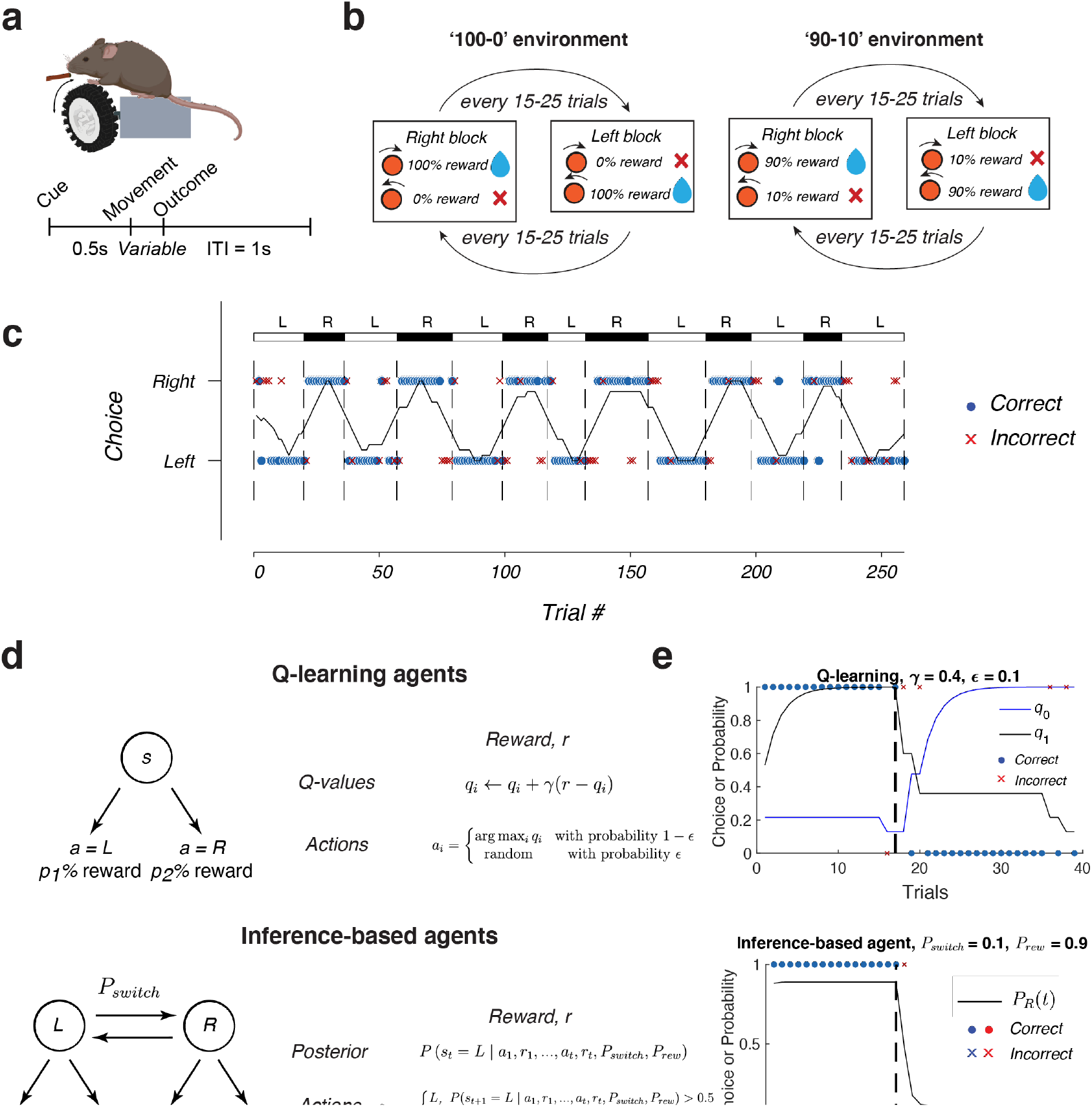
Dynamic foraging task and formulation of Q-learning and inference-based agents. a) (Top) Behavioral task setup for head-fixed mice with freely-rotating wheel. (Bottom) Timing structure for each trial, demarcating the cue, movement and outcome epochs. b) World transition models of the task. Hidden states alternated between right-states, with high reward probability for right actions, and left-states, with high reward probability for left actions. The block lengths were randomly sampled from a uniform distribution between 15-25 trials. c) Example behavioral performance of an animal in the 90-10 environment, block transitions are demarcated by vertical dashed lines. Dots and crosses represent individual trials (correct or incorrect). Black trace indicates the rolling performance of 15 trials. d) Implementation of Q-learning (top) and inference-based algorithms (bottom) for simulating choice sequences of simulated agents. e) Example behavior of simulated Q-learning (top) and inference-based (bottom). Each dot or cross represents the outcome of a single trial. In the Q-learning plot, black and blue traces represent the values of each of the two actions. In the inference-based plot, black trace represents the posterior probability of the right state *P*(*s_t_* = *R* | *a*_1_, *r*_1_,…, *a*_*t*-1_, *r*_*t*-1_).

We focused on disentangling the behavior of two classes of agents, Q-learning and inference-based agents. Q-learning is a model-free learning strategy that performs iterative value updates based on external feedback from the environment (Fig. 1d, top). In the dynamic foraging task with two options, these agents maintain two values associated with the two actions, *q_L_* for left actions and *q_R_* for right actions. On each trial, the value of the chosen action is updated toward the reward magnitude of the experienced reward, *r*, with a learning rate γ.

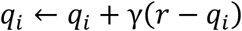

where *q_i_* represents the action value for one of the arms (*L* or *R*), *r* reflects whether the previous action was rewarded (0 or 1), and *γ* is the learning rate parameter. We additionally assumed that the agent adopts an *ε*-greedy policy. In this policy, the agent chooses the higher-valued action with probability 1 - *ε*, and chooses actions at random (with probability 50%) on a small fraction *ε* of trials. Altogether, the two free parameters, *γ* and *ε*, define a two-dimensional parameter space that captures the entire behavioral repertoire of Q-learners.

The second class of reward-based models consists of “inference-based” agents whose actions are guided by an internal model of the world. Unlike model-free agents that use the action/outcome history to directly estimate an action value for each arm, these models use the history to infer the hidden state of the environment (i.e., which side is more rewarding) and use that information to guide actions. In our task, the world model (Fig. 1) consists of two hidden states, *L* and *R*, that determine whether the “left” or “right” action is associated with higher reward probability, respectively (*P_rew_*). The evolution of these hidden states can be approximated by a Markov process with probability *P_switch_* of switching states and 1 – *P_switch_* for remaining in the same state on each trial. Given this model and observed outcomes, the ideal observer can perform Bayesian updates to keep track of the posterior distribution of the two states (see update equations in *Methods*).

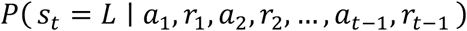

On each trial, the agent uses the posterior over the world states to select the action that maximizes the expected reward on that trial. The free parameters, *P_rew_* and *P_switch_*, constitute a two-dimensional parameter space that span the full behavioral repertoire of all inference-based agents with potentially wide variations in behavior along these two axes.

### Evaluation of previous approaches for dissociating model-free and inference-based behavior from dynamic foraging data

Dissociating model-free from inference-based behavior has traditionally been a difficult problem in this task domain. One challenge that analytical methods need to address is the large parameter space involved in these two very different models – model-free agents are described by the learning rates *γ* and exploration rates *ε*, while inference-based agents are specified by a combination of *P_switch_* and *P_rew_* of their internal models. Within these parameter spaces, the behavior can vary drastically from one region to another, requiring a thorough mapping of behavior in different parts of the two spaces before classification algorithms can be evaluated.

Due to this large size of the parameter spaces, it might not be feasible to distinguish model-free from inference-based behavior using a single behavioral metric, as previous studies have done^17,18,35^. For example, consider the use of a previously proposed feature, denoted by *ρ*, that takes into account the correlation between the number of errors in block *t* – 1, and the number of rewards in block *t*^17^. For a Q-learning agent with a low learning rate (agent denoted by blue X in Fig. 2a,d), *ρ* will be positive. This reflects the underlying slow value accumulation, such that the more rewards are experienced in the previous block, the more errors are needed in the next block to make a behavioral switch happen. On the other hand, for an inference-based agent with *P_rew_* = 0.1 and *P_switch_* = 0.7 (black X in Fig. 2b,d), the inference process is independent of the number of rewards experienced in the previous block. Thus, *ρ* is close to 0. Hence, *ρ* is a reliable metric for distinguishing the behavior of these two agents. However, this metric is insufficient to discriminate between other pairs of agents from other parts of the corresponding parameter spaces. For instance, *ρ* is also close to zero for a Q-learner with a high learning rate (blue * in Fig. 2c,d). Similarly, *ρ* may be positive for an inference-based agent with a different set of parameters (black * in Fig. 2c,d). In fact, the overall distribution of *ρ* over the two parameter spaces are very similar for the two types of models (Fig. 2d). Thus, dissociating model-free from inference-based behavior might require more detailed benchmarking of behavior using multiple complementary behavioral metrics.

**Figure 2.**
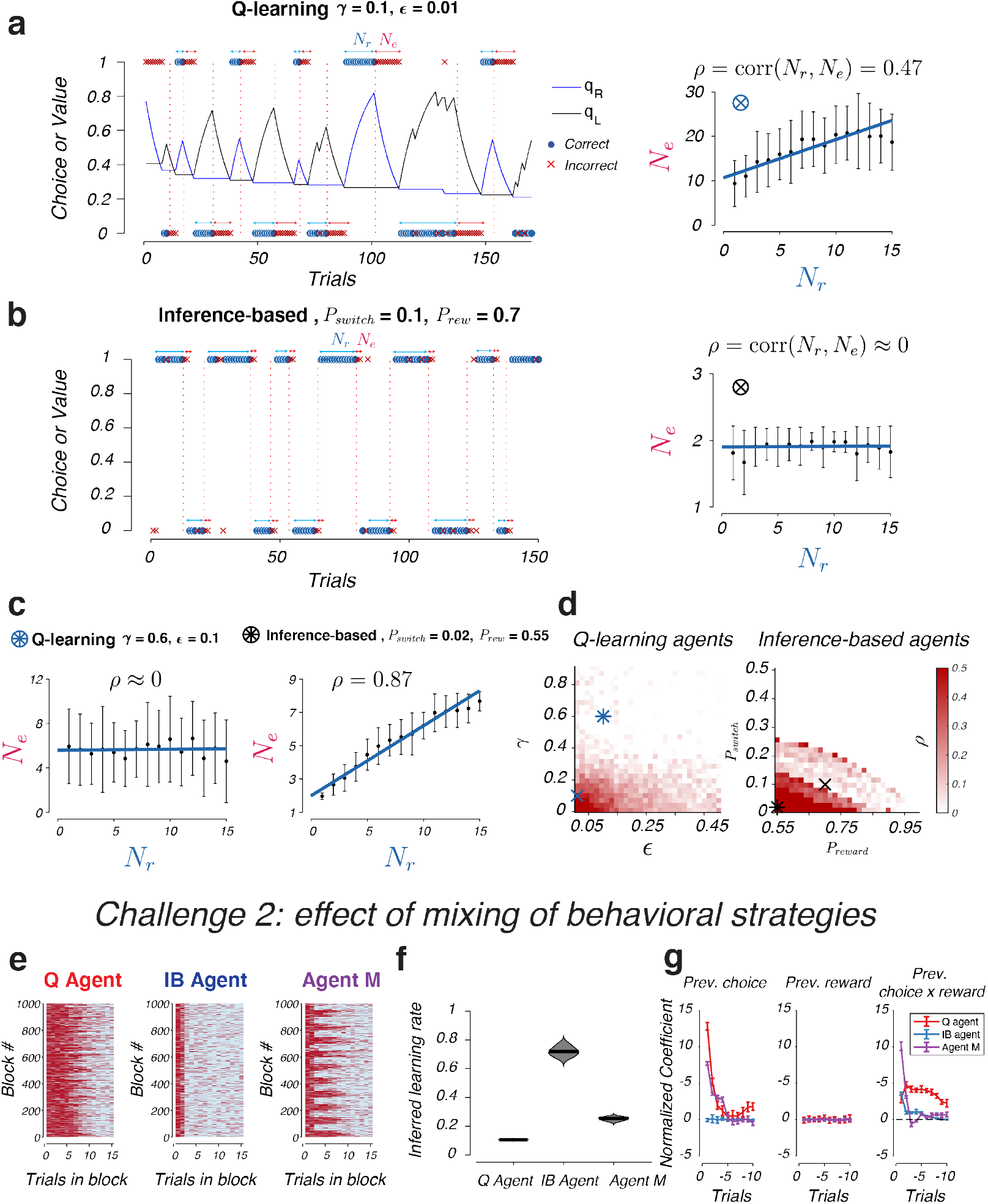
Evaluation of current analytical approaches for dissociating model-free from inference-based behavior. a) (Left) Simulation of a Q-learning agent with parameters γ = 0.1, ϵ = 0.01, illustrating the correlation between the number of errors, *N_e_*, and the number of rewards on the previous blocks, *N_r_*. (Right) Relationship between *N_e_* and *N_r_* (mean ± standard deviation, *n* = 1000 blocks across all values of *N_r_*) showing a positive correlation between the two quantities, ρ = 0.47. b) Same as a, but for an inference-based agent with parameters *P_switch_ =* 0.1, *P_rew_* = 0.7. Here, no correlation between *N_e_* and *N_r_* was seen. c) Same as a-b, but for two other Q-learning and inference-based agents that show opposite effects of *ρ*. d) Map of the values of *ρ* across the Q-learning and inference-based parameter spaces. Blue X: agent in a, black X: agent in b, blue *: Q-learning agent in c, black *: inference-based agent in c. e) Strategies of three agents over 1000 blocks of trials in the dynamic foraging task, a Q-learning agent (left), an inference-based agent (middle), and agent M (right) which mixed between the two strategies. f) Inferred learning rate by fitting the behavior of the three agents in (e) to a reinforcement learning model. Middle line represents the median (*n* = 10 repetitions). g) Logistic regression model coefficients (mean ± standard deviation, *n* = 10 repetitions) fitted on the behavioral choices of the three agents, with regressors representing previous choice, previous reward and previous choice x previous reward.

Another analytical challenge for understanding dynamic foraging behavior arises when agents mix between multiple strategies in a single behavioral session. This poses a problem for current analytical techniques such as logistic regression^15,19,24^ or reinforcement learning models^7,22,23,36,37^ which assume that the behavioral strategy is stationary within individual sessions. Although these methods work well when the agent uses a single strategy with a fixed set of parameters, they can provide erroneous estimates in scenarios of mixed strategies. To investigate the nature of such errors, we confronted models that assume stationary behavior with data generated by agents that adopt a mixture of strategies.

We simulated three agents that perform a value-guided task in a 90-10 environment (Fig. 2e). The first agent was a Q-learning agent, the second was an inference-based agent, and the third (“agent M”) mixes equally between the two strategies (see *Methods*). Both logistic regression and reinforcement learning models gave inaccurate estimates for the parameters that underlie the behavior of agent M. The learning rate inferred by the reinforcement learning model was intermediate between the two learning modes that make up agent M’s strategy (Fig. 2f).

More problematic was the result of the logistic regression model. While the inference-based agent showed no dependence on previous choice and the Q-learning agent showed positive coefficients of previous choice regressors, agent M’s dependence on previous choice was intermediate between the two agents (Fig. 2g, left panel). The coefficients for the interaction terms of agent M (previous choice x previous rewards) also showed a different pattern from either the inference-based or the Q-learning agent. Agent M’s interaction terms were higher in magnitude for the *t* – 1 trial than both the Q-learning and inference-based agents (Fig. 2g, right panel). The coefficients for previous reward are close to zero for all three types of agents (Fig. 2g, middle panel). Considering these results in the context of differentiating inference-based from model-free strategies, the inaccurate estimates are concerning. If an animal executes a mixture of inference-based and model-free strategies during the task, a method that relies on these estimates will fail to discriminate between the two modes and thus will be unable to discover the true underlying strategies.

### Four behavioral features to discriminate model-free from inference-based behavior

We first developed a framework for differentiating model-free from inference-based behavior in the case of a pure strategy with no mixing. To quantify the agent’s behavior during block transitions, we computed four features of the “transition function” that describes the dynamics of action switching of the agents in response to an uncued change in the external reward contingency (Fig. 3a). This function is a sigmoidal curve parameterized by three parameters, the switch offset, *s*, the slope α, and the lapse *ε* which represents the exploration rate of the agent in the environment. The fourth parameter is the foraging efficiency *E*, which is the fraction of rewarded choices of the agent over the whole session. In the limit of large number of blocks, this fraction is reflected by the area under the curve of the choice transition function. Either a decrease in offset, an increase in slope or a decrease in exploration would lead to an increase in the foraging efficiency.

**Figure 3.**
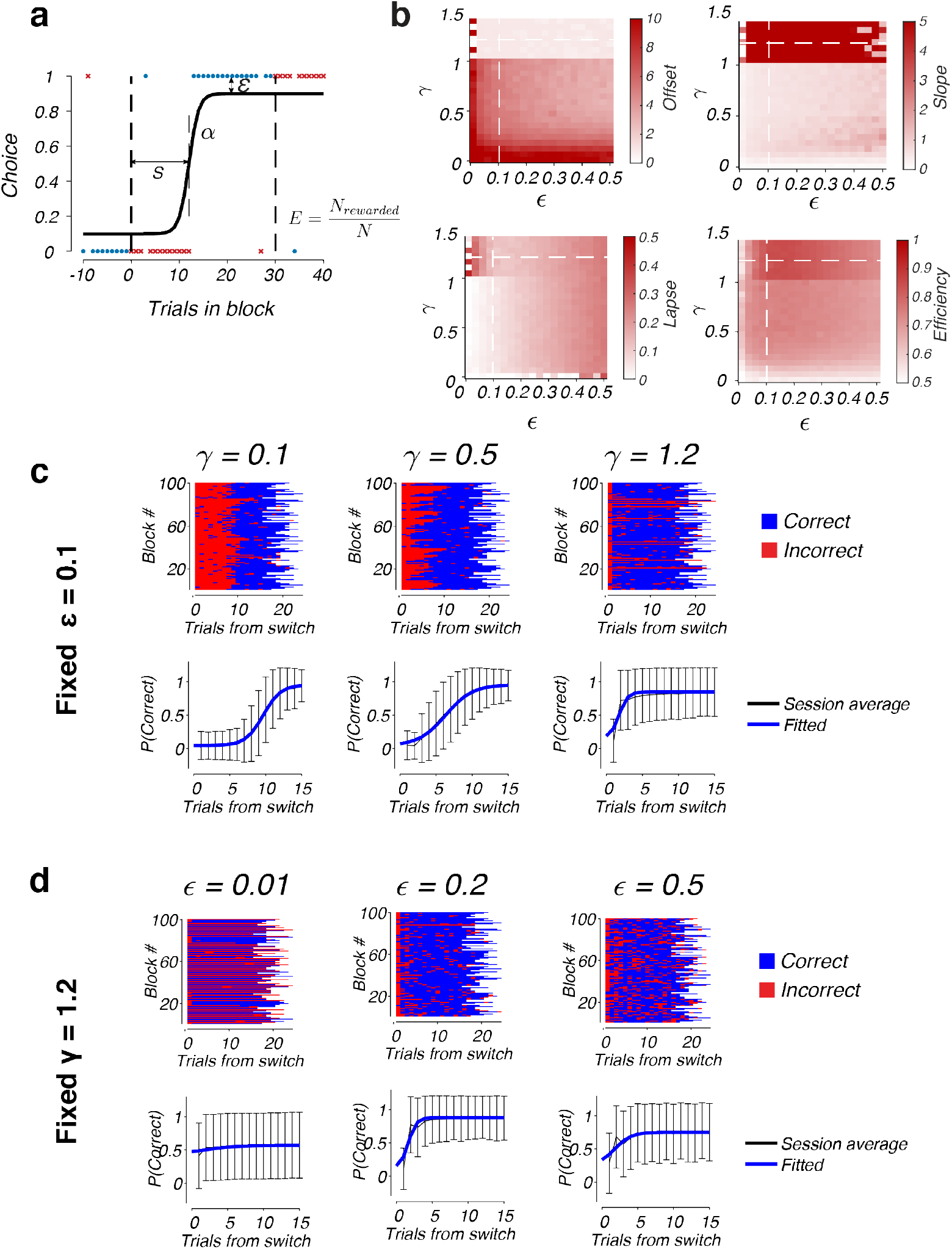
Behavioral metrics of Q-learning agents. a) Illustration of the sigmoidal transition function with four parameters: switch delay *s*, switch slope *α*, lapse *ε*, and overall foraging efficiency *E*. b) Behavior metrics for Q-learning agents in a 100-0 environment. We simulated the behavior of 25 x 20 Q-learning agents with different values of the learning rate *γ* and exploration parameter *ε*, and measured the four behavioral features for each agent by fitting the average transition function over 1000 blocks to a sigmoidal function. c) Example behavior of three Q-learning agents with a fixed ϵ = 0.1 and varying learning rate γ. Top row shows the behavior of each agent over 100 blocks (each row represents the outcomes of all the trials within a single block, red: incorrect choice, blue: correct choice). Bottom row shows the average transition function (black curve, mean ± standard deviation, *n* = 1000 blocks), and the fitted sigmoid (blue curve). d) Same as c, but for three Q-learning agents with fixed γ = 1.2 and varying *ϵ*.

We hypothesized that together, the combination of these four behavioral features can help discriminate different regimes of the model-free and inference-based behavioral spaces. For instance, the switch offset *s* might be immediate or delayed depending on the learning rate of Q-learning agents, or the parameters of the inference-based agent’s internal model. The slope *α* of the transition might be shallow or steep depending on the agent’s strategy. For an agent that relies on slow value integration from trial to trial, choice transitions might occur gradually, whereas for an agent that can quickly infer the underlying states using internal models, the transitions can be sharp. The degree of exploration might also be informative of the underlying strategy. For example, Q-learning agents require a non-zero rate of exploration in order to prevent them from getting stuck in sub-optimal strategies when reward contingencies need to be relearned. In contrast, inference-based agents with a model of the environment requires no exploration to discover these state changes. Finally, the overall foraging efficiency which non-linearly combines information from all three metrics, could be another metric that can distinguish efficient agents from less efficient ones. The use of these multiple features which are sensitive to different aspects of the behavior will thus help increase our ability to distinguish diverse ranges of behavior coming from different parts of the parameter spaces. Before building a decoder for behavioral strategy using inputs provided by these features, we will start with a survey of how each of the four features vary across the Q-learning and inference-based parameter spaces.

### Behavioral features of Q-learning agents

To characterize the behavior in the Q-learning space, we simulated an ensemble of agents, each with a different combination of γ and *ε*, where 0.01 ≤ γ ≤ 1.4, and 0.01 ≤ ϵ ≤ 0.5. For each parameter combination, we simulated the agent in the given environment (100-0, 90-10, 80-20 or 70-30) for 1000 blocks, with block sizes randomly sampled between 15-25 (similar to the protocol we use for rodent behavior training). We then averaged the behavior responses over all blocks to obtain the choice transition function (Fig. 3a), and performed a sigmoidal fit of this function to obtain the behavioral features s, α, and *ε*, that defined the switching dynamics for all points in the Q-learning space (Fig. 3b).

The distribution of behavioral features in the space was highly non-linear, and the features showed a variation along the two primary axes, *γ* and *ε* (Fig. 3b). When *ε* was held constant, a higher learning rate led to faster and sharper switching dynamics at the block transitions (Fig. 3b, c). For example, when *ε* was fixed at 0.1, increasing the learning rate *γ* from 0.1 to 1.2 led to faster behavioral switching (offset decreased from 8.6 to 5.3, to 0.8 trials). Notably, as we traversed the parameter boundary from *γ* < 1 to *γ* > 1, there was a sharp transition in the switch slope and switch offset. This is because in the high learning-rate regime where *γ* > 1, a single error was enough for agents to switch their actions, resulting in switch offsets that were very close to zero, and very sharp action transitions.

Along the *ε* axis, variations in these behavioral features were non-monotonic (Fig. 3d, top). When we fixed *γ* = 1.2, a low value of *ε* (such as *ε* = 0.01, Fig. 3d, left panel) often prevented Q-learning agents from switching as they failed to explore the alternative action after block transitions. This agent was not able to discover the more rewarding action, leading to an average transition function that is perfectly flat (Fig. 3d, bottom). A moderate value of *ε* (such as *ε* = 0.2, Fig. 3d, middle panel) encouraged exploration and enabled agents to discover the optimal action in order to make rapid action switches. However, when the degree of exploration became large (*ε* = 0.5, Fig. 3d, right panel), although the agents were able to switch rapidly, their noisy asymptotic behavior prevented them from fully exploiting the most rewarding action.

### Behavioral features of inference-based agents

Similar to the survey of the Q-learning landscape, we characterized the inference-based space by simulating an ensemble of inference-based agents with different combinations of *P_switch_* and *P_rew_* (with 0.01 ≤ *P_switch_* ≤ 0.45 and 0.55 ≤ *P_rew_* ≤ 0.99).

Unlike the variations seen in the Q-learning space which were mainly along the primary axes, the behavior of inference-based agents varied systematically along the diagonal axis of the parameter space (diagonal line in Fig. 4a). In the low *P_switch_* and low *P_rew_* regime (Fig. 4b, left panel), which we call the ‘stable’ regime of the state space, agents assumed an internal model where state transitions occur infrequently. This made them rather insensitive to errors and resulted in high switch offsets (switch offset = 8.4 trials for the agent with *P_switch_* = 0.01 and *P_rew_* = 0.55). In contrast, the regime where both *P_switch_* and *P_rew_* were high is called the ‘volatile’ regime (Fig. 4b, right panel). Here, agents assumed an environment with frequent state transitions and high reward probability. This volatile assumption made them more sensitive to errors, switching their choices more readily after only a few errors (switch offset = 0.96 trials for the agent with *P_switch_* = 0.45 and *P_rew_* = 0.99). In this regime, each error was more impactful to the agent’s update estimate of the current world state. The behavior in between these regimes had low exploration rates and offsets that were intermediate between the two extremes (Fig. 4b, middle panel).

**Figure 4.**
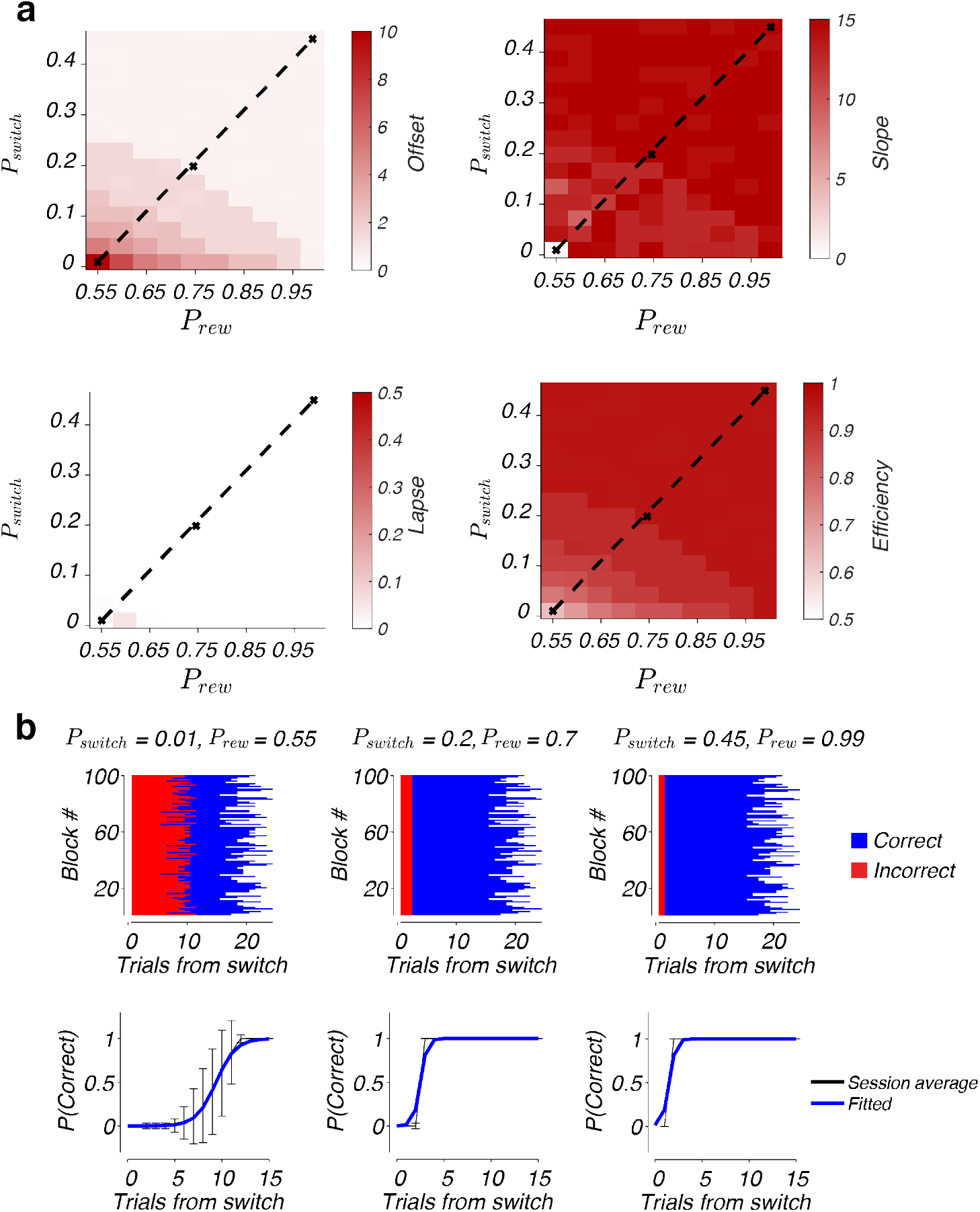
Behavioral metrics of inference-based agents. a) Behavioral features offset *s*, slope α, lapse ϵ and efficiency *E* for inference-based agents in the parameter space. Conventions are the same as Fig. 3a. b) Example behavior of three inference-based agents taken from the diagonal of the parameter space (represented by crosses in panel a plots). Conventions are as Fig. 3b,c).

One feature that distinguished inference-based agents from Q-learning agents is their lapse rates: inference-based agents tend to explore much less compared to the Q-learning agents, with lapse rates below 10% across most of the parameter space (compare Fig. 4a and Fig. 3b). This low exploration of inference-based agents can be explained by the effectiveness of the inference-based update procedure compared to the Q-learning strategy. Even for Q-learning agents with a high learning rate, a small degree of exploration is required to achieve high foraging efficiency and avoid getting stuck with low-reward actions. In contrast, Bayesian inference allows inference-based agents to infer state changes without the need to explore alternative choices. Together with the faster switch delays and sharper switch transitions, this low exploration leads to a much higher foraging efficiency than can be achieved by Q-learning agents in the uncertain worlds. Indeed, foraging efficiency was consistently above 90% for most inference-based agents, which was much higher than the maximum efficiency that can be achieved in the Q-learning parameter space (85%).

The simulation of Q-learning and inference-based agents was repeated for 90-10, 80-20, and 70-30 environments, yielding qualitatively the same trends and axes of variation among the four behavioral features in these environments (Supp. Figs. 1, 2). Thus, the qualitative trends in these features were consistent across different types of environments regardless of the level of stochasticity in the reward probability.

### Decomposition of the Q-learning and inference-based parameter spaces into sub-regimes with distinct behavioral signatures

Given the large variation of the four behavioral features across both the Q-learning and inference-based spaces, we next investigated whether the behavior of these agents naturally cluster into distinct modes that are qualitatively different from each other. To perform this analysis, we pooled the behavioral features from all Q-learning and inference-based agents in the 100-0 environment to form a 4 x 650 feature matrix, representing 4 features/agent x 650 agents (25 x 20 Q-learning and 15 x 10 inference-based agents, Fig. 5a). We applied a density-based clustering method which is well-suited for cases where the component distributions are heterogeneous and non-Gaussian^38^. The data points were first non-linearly embedded onto a two-dimensional t-SNE space, and a watershed algorithm was applied to identify borders of the embedding that separates regions of high-density point clusters. This resulted in six clusters that can be visualized on the embedding space (Fig. 5a).

**Figure 5.**
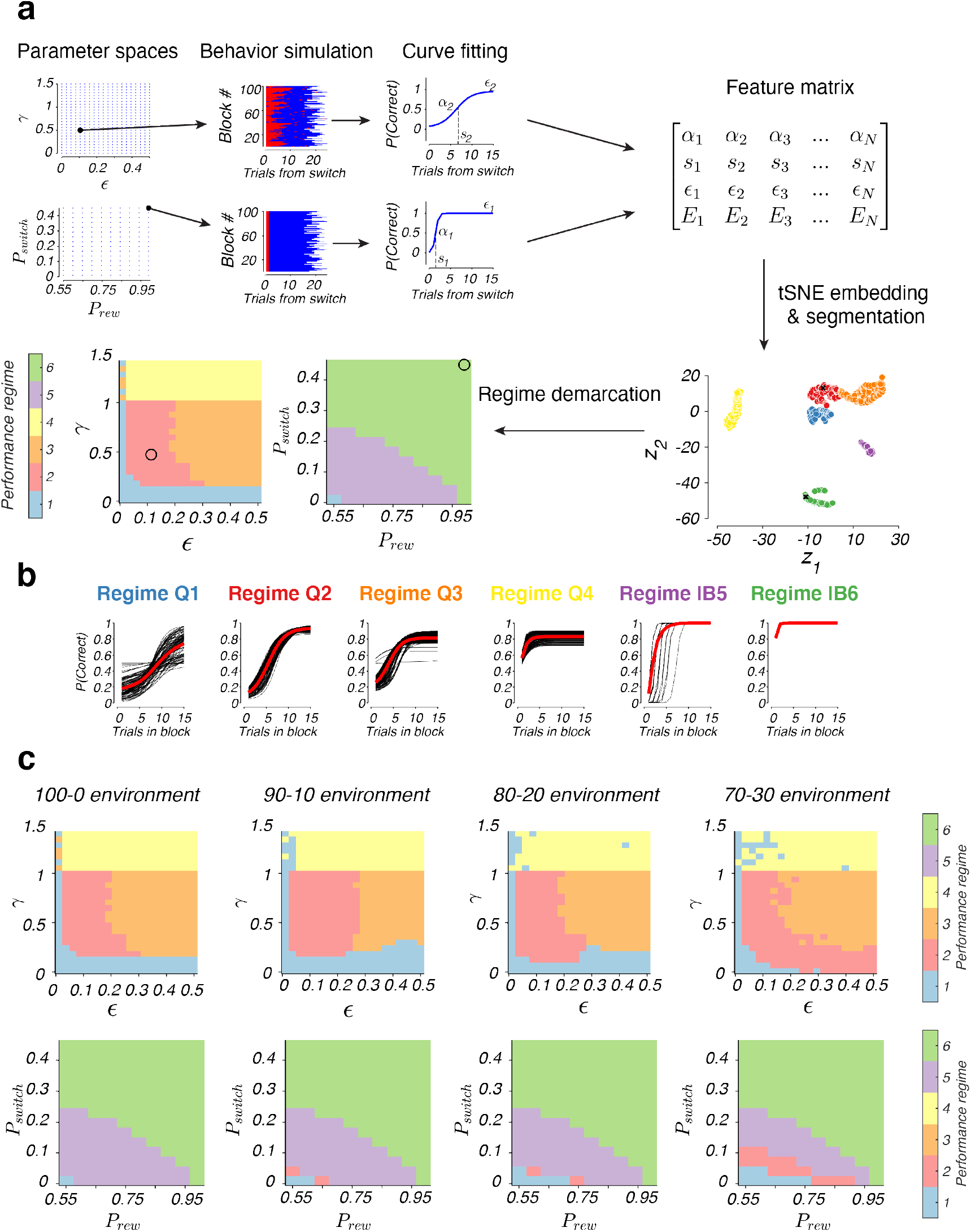
Decomposition of parameter spaces into regimes of qualitatively different behaviors. a) Method of segmentation of the parameter spaces. We performed a computational simulation of an ensemble of Q-learning and inference-based agents taken from grids that spanned the entire two spaces. For each agent, we obtained the transition function and four behavioral features characterizing the sigmoidal fit. We pooled the features of all agents into a feature matrix and applied a density-based approach to cluster these features into six regimes. We then visualized the regime identities for all points in the two parameter spaces. b) Transition functions grouped according to the behavioral regime Q1-4, IB5-6. Red trace represents the mean across all the transition functions in each group. c) Demarcation of the six regimes in the Q-learning and inference-based spaces in different types of environments (100-0, 90-10, 80-20, or 70-30).

Interestingly, when the identities of the classified points were mapped back into the parameter space that they came from, each of the six clusters corresponded to a contiguous regime in either the Q-learning or inference-based space, but not both (except for cluster 1 which was found both in large portions of the Q-learning space and a very small region of the lower left corner of the inference-based space). The first four classes were localized to regions of the Q-learning space that corresponded to low (class Q1), medium (classes Q2 and Q3) and high learning rates (class Q4), respectively. The remaining two classes were mapped to different sub-regions of the inference-based space: class IB5 resided in lower left corner of the space which corresponded to a ‘stable’ world model with low *P_switch_*; class IB6 was mapped to the complementary region, the ‘volatile’ regime where *P_switch_* and *P_rew_* are both high. The distribution of these regimes suggests a clear distinction between Q-learning and inference-based behavior, such that inference-based and Q-learning regimes are largely non-overlapping.

We verified that these regimes represented distinct modes of behavior by visualizing all the choice transition functions, grouped by the regime identity (Fig. 5b). We found qualitative differences and systematic variations across the different regime types. For example, transition functions in regime Q1 were the flattest, having shallow slopes and very late switch offset, consistent with the slow switching of Q-learning agents with low learning rates. From regime Q2 to Q4, transition functions became progressively steeper with higher slope and faster switch offsets. The average rates of exploration for all the Q-learning regimes (Q1-Q4) were all non-zero. This lapse rate was lowest for Q2 and higher in the other three regimes. In contrast, the inference-based regimes (IB5 and IB6) could be distinguished from the Q-learning clusters by lapse rates that were very close to zero. Although the behavior transitions were sharp in both regimes, they occurred at different latencies: the offset was immediate in cluster IB6 but delayed in cluster IB5, consistent with the delayed switching seen in inference-based agents with low *P_switch_* that assumed a more stable model of the world (Fig. 5b).

### Structure of behavioral features and regime demarcation in 90-10, 80-20 and 70-30 environments

So far, our clustering analysis and regime segmentation has been performed in a deterministic environment (100-0) where in each state, the reward is given with 100% probability for the high-value action and 0% probability for the low-value action. To determine how these clusters might vary in probabilistic settings, we performed the same behavior simulation and clustering procedures in 90-10, 80-20 and 70-30 environments, where rewards are given with progressively higher degrees of stochasticity. For example, in a ‘90-10’ environments, rewards are given with probability 90% on the high-valued side, and only 10% on the low-valued side. In each environment, we characterized the variations in the four behavioral features across the Q-learning and inference-based spaces (Supp. Fig. 1–2).

Our simulations revealed that the boundaries of the behavioral regimes (Q1-4 and IB5-6) were largely preserved across different environments. In all types of environments, the presence of six clusters could be confirmed when visualized in the t-SNE embeddings (Supp. Fig. 3a). Furthermore, the clusters were localized to similar regimes in the Q-learning and inference-based parameter spaces (Fig. 5c). Notably, as rewards became more unreliable (going from the 100-0 to the 70-30 environment), there was an increase in extent of overlap between Q-learning and inference-based behavior. In the 80-20 and 70-30 environments, a larger section in the lower left corner of the inference-based space was found to co-cluster with regimes Q1 and Q2 in the Q-learning space. This suggests that noisy environments, it becomes more difficult to dissociate the behavior of Q-learning agents in the Q1-Q2 regime from the behavior of inference-based agents that hold ‘stable’ internal models (the dissociability of the regimes will be further quantified by the decoding results in the next section and Fig. 6). Finally, when visualizing the behavioral transition functions of the six behavioral regimes in different types of environments, we found the same variations and patterns across the six clusters (Supp. Fig. 3b). In each environment, from regime Q1 to Q4, there was a consistent increase in the slope and a decrease in offset of the transition function. For the inference-based agents (IB5-6), we generally observed sharper transitions and faster switches compared to their Q-learning counterparts, demonstrating the usefulness of internal models in bringing about more efficient switching strategies. The IB5 cluster tended to have lower lapse rate than the IB6 cluster, and this lapse rate increased as the environment got noisier (100-0 to 70-30). As with the deterministic case, regime IB5 had a slightly delayed offset compared to IB6, as the agents’ internal belief of a stable environment made them less inclined to switch their actions as successive errors were encountered. Finally, as the level of noise increased in the environment, there was a general decrease in slope and increase in lapse rate in the transition functions for all of the six regimes.

**Figure 6.**
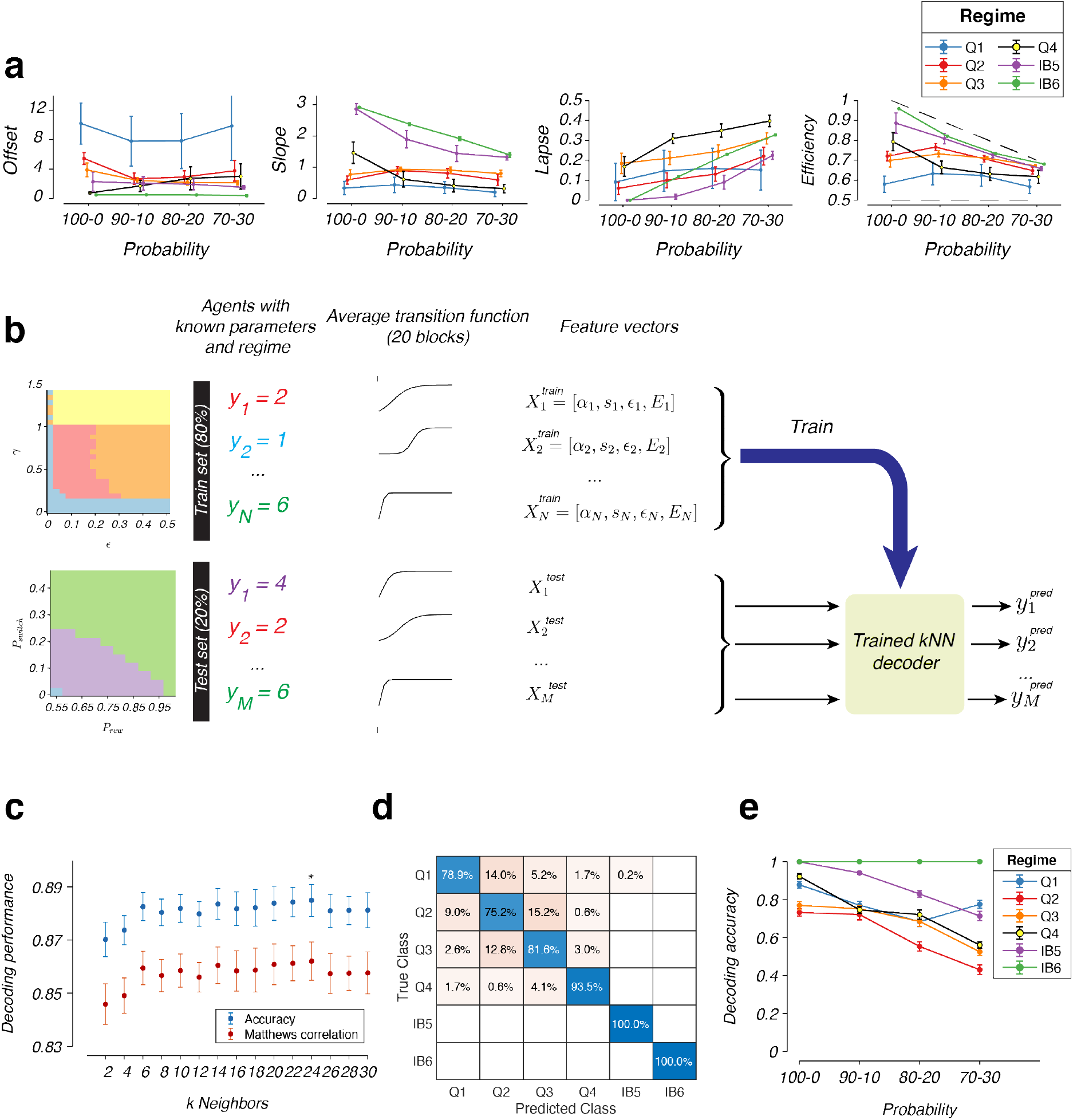
Decoding of behavioral regimes and evaluation of decoding accuracy. a) Average behavioral features (offset, slope, lapse and efficiency, mean ± standard error) of simulated agents belonging to classes 1 to 6, for the four types of environments, 100-0, 90-10, 80-20 and 70-30. In the efficiency plot (right), top dashed line represents the ideal performance, bottom dashed line represents random performance. b) Procedure for the behavioral regime decoding. c) Selection of the number of nearest neighbors, *k*, based on cross-validated decoding performance (blue, mean ± standard deviation, *n* = 20 repetitions) and Matthews Correlation Coefficient (red, mean ± standard deviation, *n* = 20 repetitions). d) Cross-validated confusion matrix for simulated behavior in the 100-0 environment. Diagonal entries show the accuracy for each respective class. e) Decoding performance (mean ± standard deviation, *n* = 20 repetitions) for the six behavioral regimes across different environments (100-0, 90-10, 80-20 and 70-30). Dashed horizontal line represents chance performance.

### Decoding of Q-learning and inference-based regime identity from behavioral data

The segregation of the Q-learning and inference-based spaces into six discrete domains suggests qualitative differences in behavior between these clusters. These differences are revealed by the features of the choice transition functions, which showed systematic variations across regime and environment types (Fig. 6a). For example, agents in regime Q1 have flattest transition functions with the highest offsets, suggesting a random mode of behavior with slow switching between the two actions. On the other hand, agents in regimes IB5 and IB6 have the lowest lapse rates and sharpest transitions (highest slopes), suggesting a mode of behavior that relies on internal models of the world to achieve the highest foraging efficiency. Altogether, these differences can be used to decode the cluster identity from the behavioral performance of animals in an experimental session. In this section, we will build and optimize these decoders, and evaluate their regime classification performance on synthetic data sets for which the ground truth is known.

The synthetic training and validation data were again obtained by computational simulations (Fig. 6b). For each agent in the Q-learning and inference-based parameter spaces (with a known regime identity according to our previous segmentation), we performed repeated simulations in 50 synthetic experimental sessions with 20 block transitions per session (chosen to resemble the number of blocks that animals typically complete in a regular training day). For each synthetic session, we averaged the behavior across all blocks to obtain the transition function, and fitted a sigmoidal curve to estimate the four features of this function. This procedure yielded a four-dimensional feature vector for each agent per session. We split this data into a training set (containing 80% of the data) and a test set (20% of the data). We trained a k-nearest neighbor (kNN) decoder on the training set to predict the behavioral regime (1 to 6), and evaluated its performance on the held-out test set. The accuracy of the decoder was measured both by the fraction of correctly labeled examples per regime, and by the Matthews Correlation Coefficient, which is a metric for evaluating the decoding performance across all six clusters (similar to the area under the ROC curve but for multi-class classifications).

We used the decoding accuracy and Matthews correlation metrics to determine the number of neighbors (*k* = 24) for optimal decoding (Fig. 6c). For the optimized decoder, the performance that could be achieved was significantly above chance for all six behavioral regimes (Fig. 6d). We found that each cluster could be decoded with higher than 75% accuracy (compared to a chance performance of 17%). Most impressively, the analysis showed that inference-based behavior (IB5-6) could be almost certainly separated from Q-learning behavior (Q1-4) (decoding performance was 99.8% for distinguishing classes IB5-6 from Q1-4 in the 100-0 environment). The decoder performed extremely well for the inference-based regimes, achieving almost perfect performance for these two clusters. The decoding accuracy was lower for classes Q1 to Q4, reflecting the higher stochasticity in these four modes due to the random exploration that is inherent in the mechanism of Q-learning agents.

We also trained separate decoders and investigated the decoding accuracy in the other three types of probabilistic environments (90-10, 80-20 and 70-30, Fig. 6e) to determine which type of environment would be the most optimal for distinguishing between the six behavioral regimes. We found that the decoding performance for the clusters dropped as the level of stochasticity increases in the environment. The decoding accuracy was consistently high and close to perfect for regime IB6, regardless of the type of environment. For each of the other five clusters, there was a drop of about 20% in decoding accuracy as we go from the 100-0 environment to the 70-30 environment. These results suggest that given our choice of behavioral features, more deterministic environments are better for distinguishing the behavior of model-free and inference-based agents, likely due to the greater separation between the behavioral features among the six types of agents (Fig. 6a).

### Session-average rodent behavior progressed through model-free regimes with increasing learning rates

The high decoding accuracy of behavioral regimes gave us more confidence to use these decoders on the experimental data that we obtained from our trained animals. We analyzed behavioral data obtained from *n* = 21 head-fixed mice that were trained on the 100-0 dynamic environment. On average, behavioral features varied systematically over time: choice transitions occurred faster (shown by the decrease in offset) and switches became sharper (shown by the increase in slope), while the lapse rate decreased with training (Fig. 7a). Although the average lapse rate decreased over time, it remained high even after 3 weeks of training (~30% on day 30), suggesting a substantial degree of exploration and indicating that not all animals transitioned to the inference-based regime at this late stage of training.

**Figure 7.**
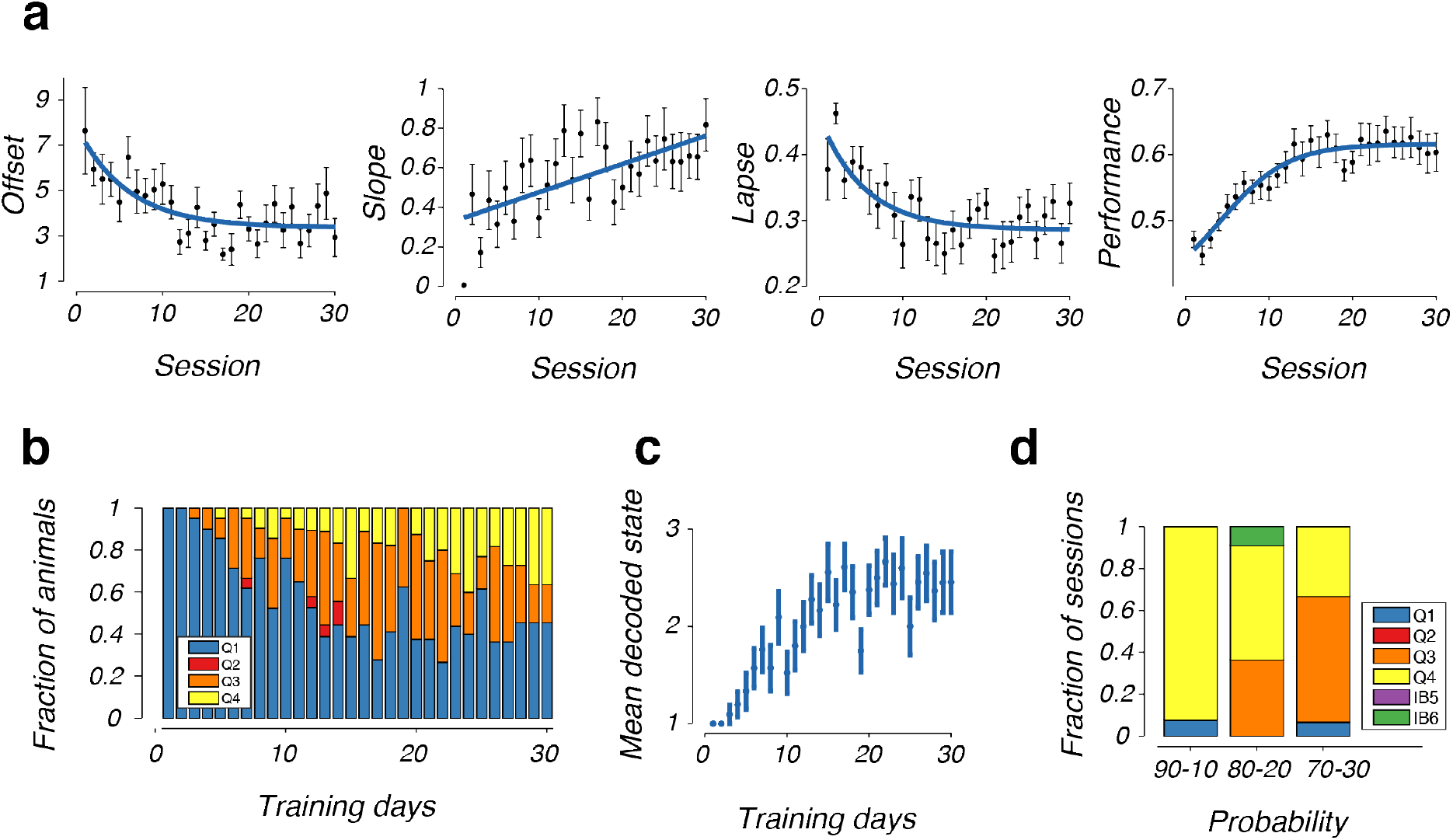
Decoding of session-averaged rodent behavior during dynamic foraging. a) Evolution of offset, slope, lapse and efficiency of rodent dynamic foraging behavior with training (mean ± standard error, *n* = 21 animals). b) Distribution of decoded state across all animals (*n* = 21) with training. c) Evolution of average decoded state across all animals (mean ± standard error, *n* = 21 animals) with training. d) Same as b, but in probabilistic environments (90-10, 80-20 and 70-30, with *n* = 6 animals). These sessions were conducted after animals became expert in the 100-0 environment.

The sharper and faster switches of trained mice in the task could be attributed to an increase in learning rate in the Q-learning mode, or a shift from the Q-learning to the inference-based decision mechanism. We dissociated these hypotheses by decoding the behavioral regime (Q1-4 or IB5-6) of each training session using the decoder that was previously trained on the synthetic data (Fig. 6). Remarkably, we found that 100% of the decoded states over the training days (across 21 animals, up to 30 training days), belonged to the Q-learning regimes, Q1-Q4 (Fig. 7b). Within these regimes, there was gradual shift toward regimes with higher learning rates. The behavior started predominantly in state Q1, and with learning, the frequency of state Q1 decreased, while states Q3 and Q4 increased in prevalence (Fig. 7b). As such, the mean decoded states across animals showed a slow increase toward higher Q-learning modes (Fig. 7c). By the end of 30 sessions, about 40% of all animals were in class Q4, and the rest were divided between regimes Q1 and Q3. There was no indication that the behavior transitioned to inference-based modes (IB5-6) in any single animal.

We also trained a subset of *n* = 6 animals on the probabilistic environments (90-10, 80-20 and 70-30). We applied decoders that are trained on synthetic data for each environment (Fig. 7d) to infer the behavioral modes for these sessions, and again found that the vast majority of these sessions were in the Q-learning regimes (Q1-Q4). Altogether, these results failed to reveal any signature of inference-based behavior from the session-averaged behavioral features of rodents. This was highly surprising, and as we noted at the start of the paper (Fig. 2d-f), could be due to the use of session-averaged statistics which can yield erroneous results by masking the use of mixtures of strategies in single sessions. In the next sections, we will tackle this challenge of analyzing mixtures of strategies by building a state-space model to quantify dynamic shifts and transitions in learning modes.

### A novel framework to quantify mixture of strategies in dynamic foraging

The absence of inference-based strategies from our previous decoding analysis was highly surprising for several reasons. First, inference-based behavior has been observed in previous studies of dynamic foraging in rodents, as well as in other complex tasks which involve multiple decision stages^17,18^. Thus, it seems unlikely that our animals are unable to develop an internal model that facilitates efficient inference in our task. Second, from our training experience, we have frequently observed expert animals making sharp switches in their actions, with some animals being able to reverse their actions after a single error after each block transition. Hence, our inability to discover inference-based behavior was suggestive of the need for a more sophisticated analysis of behavior.

One factor that might explain this result was the highly variable behavior of mice in training sessions. For example, in the same session, an individual animal might vacillate between different strategies, switching their choices immediately in some blocks, transitioning more slowly in others, and selecting choices at random toward the end of the session as they became satiated (red, green, and blue shades in Fig. 8a, respectively, for a simulated agent). These state changes pose a challenge for analysis methods which make use of session-average metrics, as highlighted by our examples in Fig. 2d-f. In our framework, each of these strategies might be governed by a separate choice transition function with varying offsets, slopes and lapse rates (sigmoidal curves in Fig. 8b). Since the session average transition function (Fig. 8a, bottom panel) is more likely to be flatter with higher lapse rate than a typical inference-based sigmoid, the average behavior will tend to look model-free, masking the inference-based strategies in some of the individual components.

**Figure 8.**
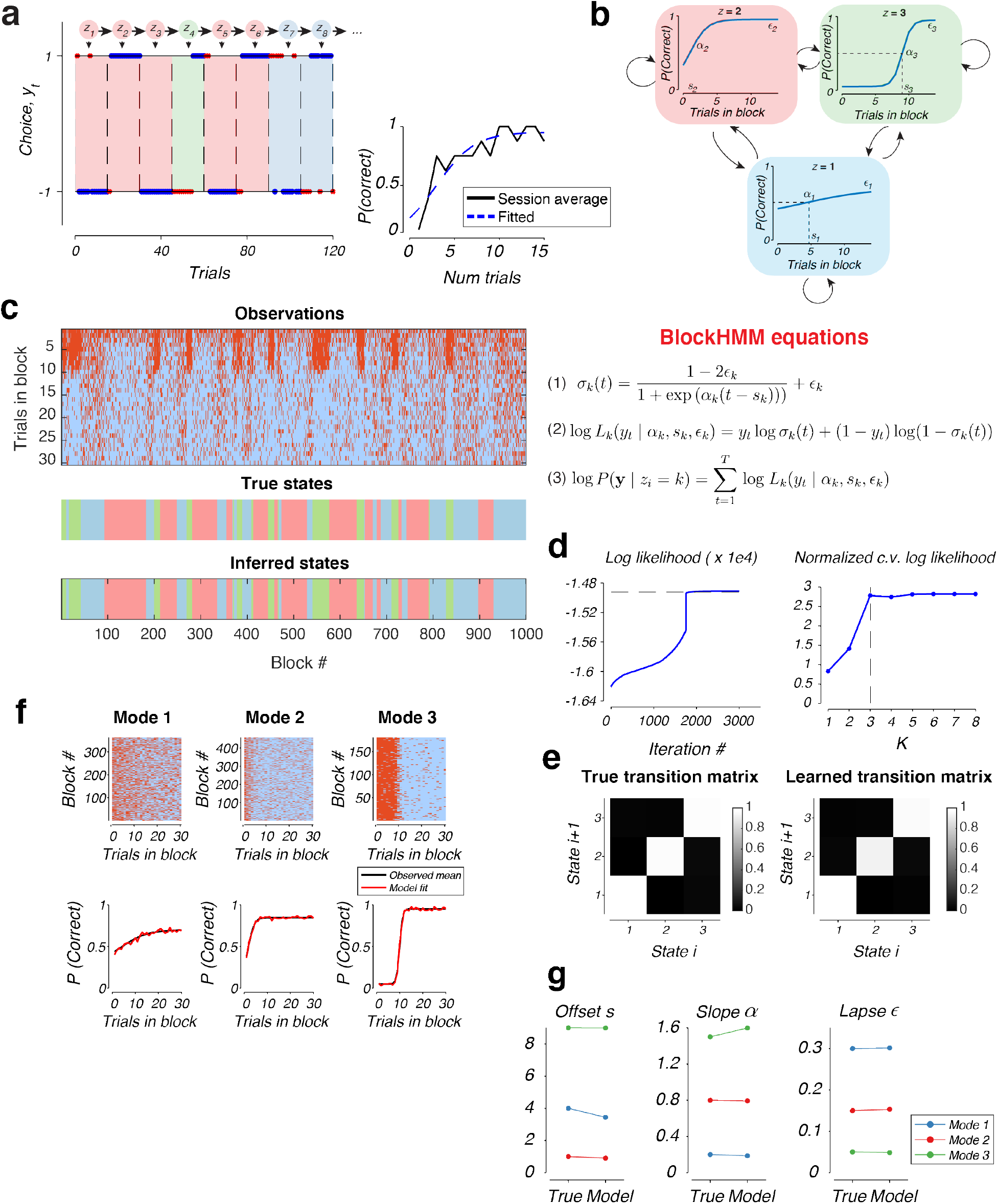
Formulation and evaluation of the blockHMM algorithm. a) Example of behavior generated by a blockHMM mixture with *K* = 3 components. The circles on top represent the underlying hidden states, *z_i_*, which evolve according to a Markov chain. Each state (shown by blue, red and green shade) follows a different set of underlying switching dynamics. Blue dots represent correct choices, red crosses represent incorrect choices. (Inset) Average transition function across all blocks of the session (black) together with the fitted sigmoidal curve (blue). b) (Top) Transition functions corresponding to each of the three hidden states, *z_i_* = 1,2,3. Each sigmoidal curve can be parameterized by three features, the slope, offset, and lapse. Arrows represent transition probabilities between the states. (Bottom) Equations of the blockHMM generative model. Each hidden state governs the choice sequence of the entire block according to the sigmoidal transitions (equations 1 and 2). The log-likelihood of the observed choices in the block is the sum of the log-likelihoods of individual trials (equation 3). c) (Top) Example behavior in 1000 blocks of trials generated by the same blockHMM mixture shown in panels a and b. Each column represents one block, with trials 1 to 30 of each block running from top to bottom. Red represents incorrect choices and blue represents correct choices. (Middle) True states that underlie the behavior shown in the top panel. (Bottom) Inferred latent states by the blockHMM fitting procedure. d) (Left) Evolution of the log-likelihood during model fitting in panel c. (Right) Dependence of cross-validated log-likelihood on the number of components, *K*. e) True and inferred transition matrices for the behavior shown in panel c. f) Grouping of blocks of trials according to the inferred state after the model fitting with *K* = 3 HMM components. (Top) Raw behavioral performance grouped by the identity of the latent state. Conventions are as Fig. 3c,d and Fig. 4b. (Bottom) Average transition function and fitted sigmoidal curve for each of the grouped blocks. g) Comparison of true and inferred parameters for the three components of the behavior shown in panel c.

The fact that individual modes of the behavior might be obscured by session-averaged measures prompted us to develop a computational tool to identify the discrete latent states that constitute the behavior of animals across their training sessions. We took advantage of recent developments of state space models that were used to infer discrete latent states from sequences of discrete or continuous variables^29,39,40^. In particular, adapting the previously developed GLM-HMM framework^29^ to the dynamic foraging setting, we assumed that each hidden state determines the parameters of a single sigmoidal transition function (offset *s*, slope α and lapse *ϵ*), which in turn determines the joint log likelihood of all the choices within each block. We named the approach “block Hidden Markov model (blockHMM)” to indicate the use of hidden states which dictate the evolution of choices throughout the block duration (Fig. 8a).

More concretely, we assumed that the choice sequences in each block *k* is governed by an underlying sigmoidal transition function *σ_k_*(t), where *t* = 0, 1, 2,… are the trial numbers within the block (Fig. 8a). These transition functions can be parameterized by the switch delay *s_k_*, slope *α_k_* and lapse rate *ϵ_k_* (Equation 1, Fig. 8b). The discrete latent states *z_i_*’s evolve from one block to the next with a Markovian property specified by the transition matrix *P*(*z*_*i*+1_ | *z_i_*) (denoted by arrows in Fig. 8a). The transition function determines the likelihood of all trials within each block (Equation 2, Fig. 8b). Finally, to fit the model, we used the EM algorithm to maximize the log-likelihood over all observed choices, which is the sum of the log-likelihoods of individual trials (Equation 3, Fig. 8b).

Our synthetic agent (Fig. 8a) was simulated according to a blockHMM process with *K* = 3 hidden states. State *z* = 1 (blue) corresponded to a random mode of behavior with a flat transition function, *z* = 2 (red) corresponded to a sigmoidal curve with a fast offset, and *z* = 3 (green) involved a sharp but delayed switching of actions. We generated the behavior of this agent over 1000 blocks (Fig. 8c), and fitted the blockHMM model to the observed choice sequences of the agent. The log-likelihood of the fit converged to the true log likelihood value (Fig. 8d, left). To determine the best number of latent states for the model, we trained the model on 80% of the blocks and evaluated the log-likelihood on the remaining 20% of the blocks. Inspecting the normalized cross-validated log-likelihood, we found that the optimal number of clusters was *K* = 3, agreeing with the ground-truth value (Fig. 8d, right). At the end of the fitting procedure, blockHMM recovered the correct transition matrix (Fig. 8e), as well as the parameters of the transition function in each mode (Fig. 8f-g). Importantly, the inferred latent states closely matched the true states that underlie the behavior (Fig. 8c, bottom panels).

### Mice use a mixture of strategies during dynamic foraging

We used the blockHMM procedure to identify the hidden states that underlie behavioral performance of our trained animals (*n* = 21). For each animal, we fit the model with the number of components, *K*, that was chosen to maximize the cross-validated log-likelihood (Supp. Fig 4, the value of *K* was also capped at a maximum value of 6 for interpretability). From the model fits, we obtained the slope, offset and lapse parameters that define each transition function. We also computed the foraging efficiency of each mode based on the performance of the animal in all of the trials in the respective states. The combination of four features per strategy were then input to our trained decoder (Fig. 6) to determine the behavioral regime (Q1-4 or IB5-6) for each of the six HMM modes (Fig. 9a). For 11/21 animals, we observed the presence of both Q-learning and inference-based regimes in the decoded HMM modes, while the rest of the animals only showed the presence of Q-learning regimes. To visualize behavior within each HMM mode, we pooled together the fitted functions from all animals (a total of 97 modes across 21 animals) and grouped them according to the decoded regime (Fig. 9b). Overall, the shape of these HMM modes closely matched the results of our regime segmentation: HMM modes that were decoded as Q1 showed delayed and gradual transitions that were close to random behavior, Q2 modes showed slow switching (with offset ~5 trials) and low exploration. Very few HMM modes were decoded to be Q3 – these modes showed similar offsets to Q2 but had higher lapse rates. Q4 modes displayed very fast switching (with offset of 1-2 trials) and a wide range of lapse rates. Importantly, blockHMM revealed the existence of a significant number of inference-based modes, which were decoded to regimes IB5-6. Consistent with our previous characterizations of these regimes, the transitions in regime IB5 occurred more slowly than IB6, and transition functions in these modes tended to have much lower lapse rates compared to the Q-learning regimes. Finally, we also recovered the state transition matrices for each animal (Supp. Fig. 6).

**Figure 9.**
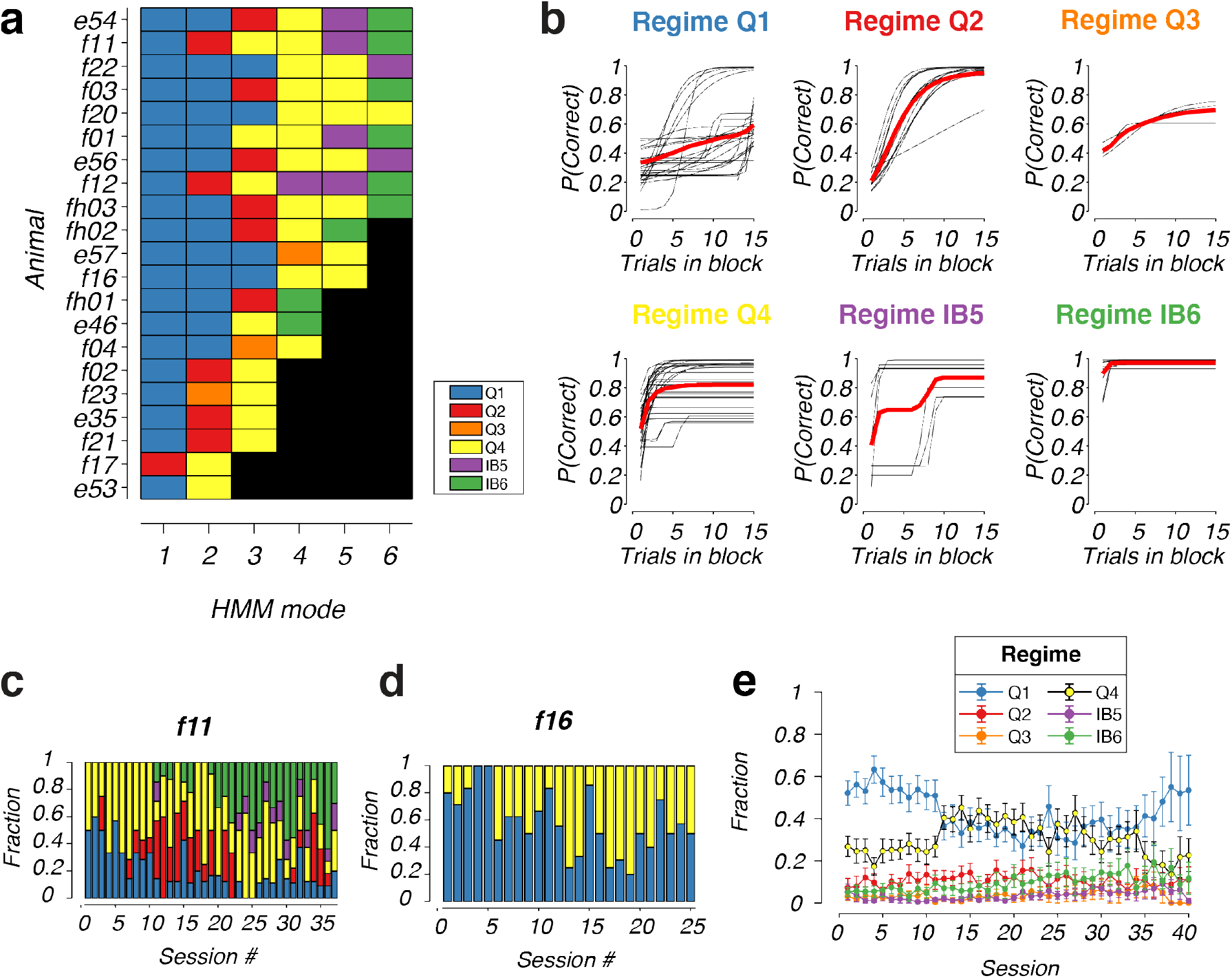
Mixture of strategies underlying rodent behavior in dynamic foraging. a) Composition of blockHMM mixtures for individual animals. Each row represents one mouse with ID shown on the left. The color of each square represents the decoded behavioral regime of each HMM mode (Q1-4, IB5-6). The number of blocks for each animal, *K*, was selected by cross-validation and are sorted here in descending order. b) Transition function of HMM modes for all animals, grouped according to the decoded behavioral regime. c) Distribution of HMM modes for an example animal, f11, across all training sessions. d) Same as c, but for another animal, f16. e) Average frequency of HMM modes for all experimental animals (mean ± standard error, *n* = 21 animals).

The model fits also allowed us to investigate the extent to which individual animals mixed between learning strategies within single training sessions. Although individual behavioral profiles were highly variable, there was a significant degree of mixing between HMM modes for all animals such that on each day, it was common to see a mixture of two or more behavioral regimes. An example animal (f11, Fig. 9c) executed an approximately equal mixture of Q1 and Q4 on its first training days. This fraction slowly shifted over time, as the prevalence of the Q1 mode decreased, while other Q-learning modes with higher learning rates (such as Q2-Q4) started to dominate. Around day 10 of training, the inference-based modes started to appear, growing in proportion until the late stages of training. However, remarkably, even in the expert stage (day 38 of training), the animal never operated fully in the inference-based regime. Instead, there remained a mixture of both inference-based and Q-learning strategies in roughly equal proportions at this stage of training. This was a common feature of many animals that managed to reach the inference-based stage (such as animal e46, e54, e56, f01, f11, f12, fh02, fh03, Supp. Fig. 5). On the other hand, a small subset of other animals, such as f16 (Fig. 9d), never reached the inference-based mode even after up to 25 days of training. The behavior of these animals primarily mixed between regimes Q1 and Q4 on each day, presumably alternating between periods of attention (high learning rate, Q4) and low attention (low learning rate, Q1).

We compared the results of our segmentation approach to previously proposed metrics to distinguish between model-free and inference-based behavior. We determined the ρ coefficient as defined in Fig. 2 and previous studies^17^, for all training sessions across our *n* = 21 animals. On average, ρ systematically shifted from a significantly positive value for the first 5 sessions (Supp. Fig. 7a, *p* < 10^-5^, Wilcoxon signed-rank test, *n* = 21 animals) to a value that is not significantly different from 0 for sessions 21-25 (Supp. Fig. 7a, *p* = 0.3, Wilcoxon signed-rank test, *n* = 21 animals). This is consistent with the previously reported trend^17^ and the average transition from model-free to inference-based modes of learning in our animals. However, the trends in *ρ* for single animals were noisy (Supp. Fig. 7b, c) which made it more challenging to distinguish model-free from inference-based behavior in single sessions. For example, although the two animals f11 and f16 (Fig. 9c,d) had qualitatively different behavioral modes as revealed by blockHMM, the evolution of the *ρ* estimates were qualitatively similar and not statistically different from sessions 21 – 25 (Supp. Fig. 7b, *ρ* = 0.8, Wilcoxon signed-rank test, *n* = 5 sessions). Moreover, for animals whose behavior primarily lie in the Q-learning regime (e53, e57, f04, f16, f20), p was not statistically different from 0 in many sessions. This discrepancy could be attributed to the level of noise in the estimates, or the fact that Q-learning agents can also have *ρ* values close to zero especially in the high-learning rate regime (Fig. 2c, d).

Across all animals, the average frequency and dominance of the HMM modes and behavioral strategies changed systematically over the course of training (Fig. 9e). On average, animals started training with a significant fraction of the Q1 mode and smaller fraction of Q4 (56% in Q1 and 24% in Q4, averaged across days 1-5). Over the course of training, the mixture of behavioral strategies slowly shifted from Q1 to Q4, such that around day 15, there is a higher fraction of Q4 than Q1 mode (39% in Q4 compared to 35% in Q1, averaged across days 16-20). This shift in composition reflects an average increase in learning rate in the Q-learning regime. At the same time, the fraction of inference-based modes, IB5 and IB6, was low at the beginning (3% in IB5 and 6% in IB6 averaged across days 1-5), but continuously increased as animals gained experience with the task (6% in IB5 and 14% in IB6 averaged across days 36-40). Notably, at the expert stage, there was a significant fraction of blocks in the inference-based mode (20% in IB5-6 combined averaged across days 36-40), but the mixture of strategies still remained with Q1 and Q4 being the primary Q-learning modes of the animals. Overall, these ubiquitous use of mixtures of strategies, which were distinctive both in naïve and expert animals, further underscore the importance of our approach to dissociate and characterize the features that constitute individual modes of behavior.

## Discussion

Model-free and inference-based strategies are the two types of models that are most often used for analysis of choice sequences in dynamic foraging experiments. Model-free constructs such as reinforcement learning models have been particularly useful when probing representation of action values in numerous brain regions^7,14,15,22,41–43^. Complementarily, inference-based models using Bayesian inferences has helped us understand the inference process that occurs in the brain from trial to trial when animals hold an internal model of the world transitions^18,26^. In the dynamic foraging task, while most studies tend to focus exclusively on one of the two model types, it has been recognized that both of these modes can co-exist in the behavior of humans and rodents, with a transition from model-free to inference-based behavior as they gain familiarity with the task^17,44^. By providing the tools to understand the difference between these two modes of behavior, our study provides a basis for comparison between these two disparate spaces of models. Our efforts are among other work of dissociating model-free from inference-based (or model-based) behavior in other task domains^45–48^. As building internal models of the world is a crucial, challenging but less understood brain function^49^, distinguishing between model-free from inference-based behavior in dynamic foraging is the first step toward an understanding of how these internal models can be acquired with learning.

Our approach builds upon previous work in this domain in several ways. First, we proposed a framework that relies on quantitative measurements of four behavioral features that characterize transitions between actions, using the concept of transition functions which had only been qualitatively characterized by other studies^19^. Our combined use of four behavior features also makes it easier to decode the behavioral strategies, as these metrics offer better coverage of the large parameter spaces involved in the two models, *γ*-*ε* for Q-learning agents, or *P_switch_* - *P_rew_* for inference-based agents. Although we have not considered other behavior features such as the probability of action switching^19,26^, similar metrics can be incorporated in the same framework to potentially improve the decodability of strategies even further. In general, the use of multiple features would help maximize the discriminability between the two types of behavior in the high-dimensional feature space. This offers an improvement from previous attempts which use a single parameter to distinguish between the two modes of learning. For example, we showed that *ρ* by itself is insufficient to distinguish model-free from inference-based behavior for certain pairs of agents^17^. In the same way, this problem also applies to other single metrics such as transition slope^18^ or offset^35^ which have been used in previous studies. Our approach also differs from previous attempts using data-driven methods^27^ to predict the choice of animals and agents on individual trials. We instead try to estimate a set of aggregate behavioral metrics such as the switch offset and lapse rate to decode the behavioral regimes of different agents. Since our focus is to predict the behavioral class rather than the choice on single trials, this allows us to gain statistical power and robustness as these aggregate measures can be estimated more accurately from the performance of the animals over multiple blocks of trials, in comparison to single-trial choice prediction which can be difficult due to the presence of noise in the choice sequences^50^.

Among the four features we investigated, the variation in lapse rate during training was particularly noteworthy. We found that there was a high lapse rate in our experimental animals, even in the deterministic environment where the reward contingency should be straightforward to learn and acquire. On average in this easiest task condition, the lapse rate of animals even on day 30 of training was close to 30%. In our Q-learning model, this lapse rate could be accounted for by a high value of *ε* which leads to a high degree of exploration of the animals. This high rate of exploration would agree with previous studies of mice in a maze^51^, open-field^52^ or head-fixed^9^, which found a tendency for mice to explore their environments, presumably to gain information about unknown events or contingencies^53,54^. Although exploration is the most direct explanation for the high lapse rate, we cannot rule out the possibility that the high lapse rate could be caused to other factors such as inattention, motor errors, or incomplete knowledge of the task^29,55–57^, which similarly affect the interpretation of lapse rates in sensory-guided behavioral paradigms^9,29^.

Together, the four behavioral features of the transition dynamics, the switch offset, slope, lapse and efficiency, provide a basis for reliably classify the behavior of different Q-learning and inference-based agents into one of six distinct clusters that show qualitatively different behavioral phenotypes. Remarkably, each of these two parameter spaces can be further segmented into smaller subdomains, thus highlighting the heterogeneity of behavior within these two classes of strategies. We found that the Q-learning space can be divided into four clusters, Q1-Q4, that broadly correspond to different learning rates. Q1 is a low-learning rate regime where the behavior is close to random on most of the block, Q2,3 have moderate learning rates where slow block transitions occur, while Q4 is a high-learning rate regime where the behavior transitions are fast, but foraging efficiency can be strongly dependent on the degree of exploration, highlighting the well-known exploration-exploitation trade-off in reinforcement learning^58,59^. In this regime, too low exploration risks getting the agent stuck a sub-optimal choice during block transitions, while too high exploration results in a failure to maximize received rewards. The types of behavior for model-free agents might be even more complex when alternative schemes for exploration, such as soft-max, UCB-1 or pursuit^59^, are considered. Interestingly, in our characterization, the difference between lapse rates turned out to be an important criterion for distinguishing model-free from inference-based behavior, especially in deterministic (100-0) environments. Here, model-free clusters (Q1-Q4) tend to have significant, non-zero rates of exploration, while inference-based clusters (IB5-6) has a lapse rate that is very close to zero. This suggests that the lapse rate can serve as an additional discriminator for the two types of models, in addition to other metrics that have been considered by previous studies^17,18^.

The ground-truth parameters used in our simulations also allowed us to evaluate the reliability of decoding model-free from inference-based behavior in different types of environments. We found that decoding accuracy was highest in the deterministic (100-0) environment and slowly degrades for more stochastic environments (going from 90-10 to 80-20 and 70-30). This degradation arises because in probabilistic environments, inference-based and model-free transition functions become more similar. In such noisy environments, an efficient inference-based procedure might still give rise to slow and delayed switching since in these environments, the rewards received are rather uninformative of the current state of the world. The lapse rates of inference-based agents also become non-zero in this unreliable condition which makes it difficult to distinguish between the effect of *ε*-greedy exploration in Q-learning agents. On the other hand, in the deterministic, 100-0 environment, a failure to fully exploit an action after switching must be attributed to exploration, allowing an accurate detection of exploration states which imply a Q-learning behavior. The decoding accuracy of behavioral strategies thus establishes a baseline evaluation of our ability to distinguish model-free from inference-based behavior in high-noise environments.

The second major contribution of this work is the development of a state-space model, blockHMM, which allows us to segment of behavior during the session into blocks of trials that are governed by different underlying states. Our work adds to the existing body of literature for quantifying mixtures of strategies in reward-guided contexts which revealed interacting components of behavior involving reinforcement learning, working memory, episodic memory or the interaction between model-free and model-based systems^27,60^. To tackle challenges faced by models that assume stationarity of behavior (Fig. 2e-g), our model takes inspiration from recent modeling approaches which are used to infer discrete latent states that underlie neural dynamics^39^, natural behavior^40^, and behavior in decision-making tasks^28,29^. In particular, we adapted the recent GLM-HMM framework^29^, where discrete hidden states determine the coefficients of a generalized linear model (GLM) which specifies how the decision of the animal depends on external trial variables. While the latent states in this approach are updated from trial to trial, latent states in the blockHMM framework govern the choice selection across entire blocks, and are only updated at the boundaries of block transitions. Each state involves a separate sigmoidal transition function parameterized by the slope, offset and switch. By pooling the behavior across different sessions, blockHMM bootstraps from the large number of blocks across multiple sessions to estimate these state-specific parameters. As these are the same parameters that are used for decoding Q-learning or inference-based regimes, this allows us to recover the behavioral regime (Q1-4 or IB5-6) that corresponds to each state. We performed a cross-validation analysis to determine the number of states, *K*, that best describe the behavior of each animal, ensuring that these modes are meaningful units of behavioral states and not arbitrary noise patterns that are fit by the model.

Our results uncover a remarkable diversity of behavior across the 21 animals that were trained in the task. This diversity is demonstrated by different number of HMM modes, *K*, the composition of the modes (Fig. 9a), the shapes of the transition function of each mode (Fig. 9b), the transition probabilities (Supp. Fig. 5), as well as the evolution of the mixture composition throughout the course of training (Supp. Fig. 4). We found only 11/21 of our animals transitioned to an inference-based mode of learning, while the rest of the animals remained in the Q-learning modes. This might explain why some previous studies might not observe efficient inference-based behavior of rodents during behavioral switching^19^, since a large fraction of animals might have failed to transition to this regime.

Not only is the behavior variable across animals, but it can also be highly dynamic within a session. We found that rodents frequently employ a mixture of strategies, mixing between periods of random behavior, Q-learning and inference-based behavior even at the expert stage after being exposed to the task for many weeks. This is so even for the easiest reward contingency (100-0 environment) where the optimal decision is simple – the animal only needs to make a switch each time a single error is encountered. Although we might expect rodents to be able to quickly figure out this task and become fully committed to the inference-based strategy, this was not the case. Instead, the frequent switches between behavioral states is representative of rodent behavior and agrees with many other studies of a diverse array of tasks^28,29^. This feature of rodent behavior once again highlights the need for powerful analytical methods that can infer hidden behavioral states that govern behavior, since these types of models allow a finer scale resolution when dissecting the behavioral circuits.

Overall, our study lays the foundation for future analyses and investigations into the neural basis of model-free and inference-based behavior, and calls for a focus on the problem of state segmentation in rodent behavioral studies. An interesting question that is raised by our characterizations is how internal models are acquired during the task, and the factors that affect the evolution of parameters of these internal models. The methods developed in the paper can be leveraged in investigations of the neural mechanisms that govern these distinct modes, as well as the plasticity of these circuits during the transition between model-free and inference-based behavior. The state segmentation approach will also be a valuable tool for perturbation experiments, with the power to reveal shifts in composition, order or transition probabilities between these modes, thus augmenting existing methods for a much richer and complete view of rodent behavior during dynamic foraging.

## Materials and Methods

### Animals

All experimental procedures performed on mice were approved by the Massachusetts Institute of Technology Animal Care and Use Committee. Mice were housed on a 12 h light/dark cycle with temperature (70 ± 2 °F) and humidity (30–70%) control. Animals were group-housed before surgery and singly housed afterwards. Adult mice (2-6 months) of either sex were used for these studies. In addition to wild-type mice (C57BL/6J), the following transgenic lines were used: Ai184D (B6.Cg-Igs7tm148.1(tetO-GCaMP6f,CAG-tTA2)Hze/J), Jackson #030328; Ai162D (B6.Cg-Igs7tm162.1(tetO-GCaMP6s,CAG-tTA2)Hze/J), Jackson #031562; B6.129(Cg)-Slc6a4tm1(cre)Xz/J, Jackson #014554.

### Surgical procedures

Surgeries were performed under isoflurane anesthesia (3–4% induction, 1–2.5% maintenance). Animals were given analgesia (slow release buprenex 0.1 mg/kg and Meloxicam 0.1 mg/kg) before surgery and their recovery was monitored daily for 72 h. Once anesthetized, animals were fixed in a stereotaxic frame. The scalp was sterilized with betadine and ethanol. The skull was attached to a stainless-steel custom-designed headplate (eMachines.com) using Metabond. Animals were allowed to recover for at least 5 days before commencing water restriction for behavioral experiments.

### Behavioral apparatus and task training

The training apparatus and software for running the experiments were adapted from the Rigbox framework for psychophysics experiments in rodents^61,62^. Mice were head-fixed on the platform (built from Thorlabs hardware parts) and their body placed in a polypropylene tube to limit the amount of movement and increase comfort. Their paws rested on a vertical Lego wheel (radius 31 mm) which was coupled to a rotary encoder (E6B2-CWZ6C, Omron), which provided input to a data acquisition board (BNC-2110, National Instruments). The data acquisition board also provided outputs to a solenoid valve (#003-0137-900, Parker) which controlled the water reward delivery.

After mice recovered from surgery, they were placed under water restriction for 1 week, with daily water given by HydroGel (Clear H_2_O). The initial amount of HydroGel was equivalent to 2mL of water a day, and this decreased gradually until mice received an amount equivalent to 40 mL/kg each day. Mice were weighed weekly and monitored signs of distress during the course of training. In the case of substantial weight loss (>10% loss weekly) or decrease in body condition score, the restricted water amount was increased accordingly. Mice were handled daily during the initial 1-week water restriction period for ~10 minutes each day. They were then allowed to explore the apparatus and given water manually by a syringe on the platform. If mice did not receive their daily water amounts during training, they were given the remaining amount as hydrogel (Clear H_2_O) in their home cage.

When mice were comfortable with the setup, they were head-fixed on the platform and given small water rewards of 4 μL from a lick spout every 10 seconds, for a total duration of 10 minutes. This duration was increased to 20 minutes, and 40 minutes on the two subsequent days. The wheel was fixed during this protocol. On the next day, mice were trained on the *movementWorld* protocol, with the wheel freely moving. Here, each trial was signal with an auditory tone (0.5s, 5 kHz), following which movements in any direction crossing the movement threshold of 8.1° rotation were rewarded with 4 μL of water. Mice then had to remain stationary for 0.5 s before the next trial starts. This discouraged a strategy of continuous rotation of the wheel.

After mice became comfortable with this stage and consistently obtained at least 0.6 mL of water each session, they were taken to the final task stage, *blockWorldRolling*. Each trial began with an auditory tone (0.5s, 5 kHz). During a delay period of 0.5 s from the trial tone onset, movements of the wheel were discounted. After this window, the movement period started, where movements of the wheel past a specified threshold were recorded. The threshold was fixed at 8.1° in the first session of *blockWorldRolling* and subsequently increased to 9.5°, and 10.8° on the next days. The trials were grouped into blocks of trials of 15-25 trials, with lengths of the blocks sampled uniformly at random. The blocks alternated between the “left” and “right” state. In the “left” state, left wheel turns were rewarded with probability 100% and right wheel turns were not rewarded. In the “right” state, right wheel turns were rewarded with probability 100% and left wheel turns were not rewarded. If mice made the correct movement, they were given a 4 μL water reward. For unrewarded trials, a white noise sound was played for 0.5 s, followed by a time-out of 1 s. After the trial feedback was given, an inter-trial interval (ITI) of 0.5 s elapsed before the next trial started. The ITI was gradually increased to 1 s once animals performed well in the task. If mice didn’t make a choice within 20 seconds, the trial was aborted, signaled by a white noise and 1-s time-out period (similar to an error trial). After the length of the block has passed, if the rolling performance of the animal in the last 15 trials was above 75%, the state of the block would flip and the next block continued. Otherwise, the block continued until the rolling performance in the last 15 trials in the block passed 75%.

For *n* = 6 animals (F11, F12, F16, F17, F20, F21), after becoming expert in the 100-0 environment, we continued training them in successively more volatile environments. Each animal was trained in 2-3 sessions in the 90-10 environment, followed by 2-3 sessions in each of the 80-20, and 70-30 environments. The example behavior in Fig. 1c was for animal F11 on a 90-10 environment.

### Simulated environment

We simulated an artificial environment that alternates between two states, “left” and “right”, in blocks of trials. The first block was chosen at random to be in the “left” or “right” state, and the state identity flipped for each subsequent block. At the start of each block, we determined the number of trials in the blocks, *N*, by sampling an integer at random in the range [15, 25]. We then simulated *N* trials in the block. In each trial, the agent selected an action (see “Simulation of Q-learning agents” and “Simulation of inference-based agents” for details below) and received feedback from the environment. If the block was in the “left” state, left actions yielded reward with probability of *p* and right actions yielded reward with probability of 1 – *p*. Conversely, if the block was in the “right” state, left actions yielded reward with probability of 1 – *p* and right actions yielded reward with probability of *p*. We considered four different environments with *p* = 1.0, 0.9. 0.8 and 0.7, which we called 100-0, 90-10, 80-20 and 70-30, respectively.

### Simulation of Q-learning agents

Each Q-learning agent was defined by two parameters, the learning rate γ and exploration rate ϵ. For our simulations, we simulated a 25 x 20 grid of parameters within the range 0.01 ≤ γ ≤ 1.4, and 0.01 ≤ ϵ ≤ 0.5.

On each trial, the Q-learning agent implemented a Q-value update and selected actions with an ϵ-greedy policy. The agent maintained two values associated with the two actions, *q_L_* for left actions and *q_R_* for right actions. We initialized *q_L_* = *q_R_* = 0.5. On each trial, the agent updated these values according to

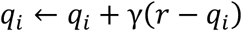

where *r* is the feedback of the trial (*r* = 1 for rewarded actions and *r* = 0 for non-rewarded actions). The Q-learner chose the higher-valued action with probability 1 - *ε*, and selected actions at random (with probability 50% for each choice) on a small fraction *ε* of trials.

### Simulation of inference-based agents

Each inference-based agent held an internal model which consisted of two hidden states, *L* and *R*, that corresponded to the unobserved hidden states, “left” or “right”, of the environment. The internal model was defined by two parameters, *P_switch_* and *P_rew_* according to

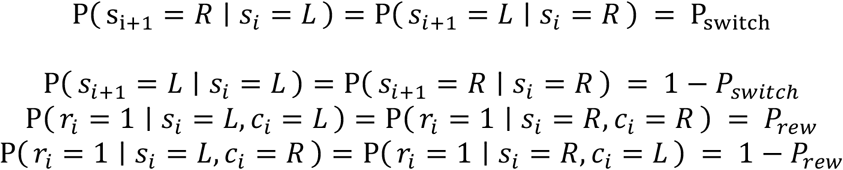

where *s_i_* refers to the hidden state on trial I and *c*_*i* refers to the choice on trial i.

That is, the evolution of the hidden states followed a Markov process with probability *P_switch_* of switching states and 1 – *P_switch_* for remaining in the same state on each trial. For our simulations, we simulated a 15 x 10 grid of parameters within the range 0.01 ≤ *P_switch_* ≤ 0.45, and 0.55 ≤ *p_rew_* ≤ 0.99.

We derived a recursive update for the agent’s posterior belief about the current world state, given previous choices and feedback. Let P_L_(t) = (*s_t_* = *L* | *c*_1_,*r*_1_,*c*_2_,*r*_2_,…,*C*_*t*-1_,*r*_*t*-1_) and *P_R_*(*t*) = (*s_t_* = *R* | *c*_1_,*r*_1_,*c*_2_,*r*_2_,…,*C*_*t*–1_,*r*_*t*–1_). Then

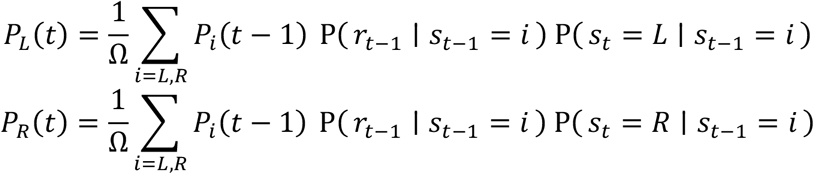

where Ω is a normalization factor to ensure *P_L_* (*t*) + *P_R_* (*t*) = 1.

We initialized *P_L_*(0) = *P_L_*(0) = 0.5. On each trial, the agent selected the left action if *P_L_*(*t*) > 0.5, the right action if *P_L_* (*t*) < 0.5, and acted randomly otherwise.

### Evaluation of previous metrics and approaches

For a given agent, the ρ metric is defined as follows. For each block transition, we counted the number of consecutive rewards that take place before the block transition, *N_r_*, and the number of consecutive errors that take place immediately after the block transition, *N_e_*. We defined ρ to be the Pearson correlation coefficient between *N_r_* and *N_e_* across all the blocks in the session. To minimize the effect of outliers, we only considered blocks where *N_r_* ≤ 15.

The Q-learning and inference-based agents in Fig. 2a-d were simulated in a 90-10 environment, where the block lengths ranged from 5 to 40. The block lengths were sampled as follows. The minimum possible block length was 5 trials, and each subsequent trial where the agent chose the high-reward side, there was a 10% probability of switching states. The block also automatically switched after 40 trials had elapsed.

The Q-learning agent in Fig. 2e was simulated with γ = 0.1 and ϵ = 0.1. The inference-based agent in Fig. 2e was simulated with *P_rew_* = 0.7 and *P_switch_* = 0.2. Each agent was simulated for 10 total sessions, each lasting 1000 blocks. For agent M, we used a mixture of strategies: we alternated between the Q-learner’s strategy for 50 blocks and the inference-based agent’s strategy for 50 blocks, and kept alternating between these modes until the agent has executed 1000 blocks in total. This was repeated for 10 total sessions (similar to the Q-learning and inference-based agents) to obtain error bars for the parameter estimates.

To infer the learning rates in a traditional reinforcement learning framework (Fig. 2f), we fit a reinforcement learning model with three parameters, learning rate γ, inverse temperature β, and bias *b*, to the sequence of choices and feedback of the agent. We assumed the agent maintained Q-values for the left and right action and use the same update rules as described in “Simulation of Q-learning agents”. Given Q-values *q_L_* and *q_R_*, the likelihood of selecting an action is given by

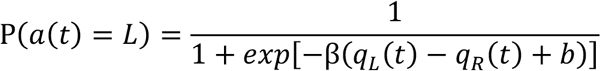

We jointly fit γ, β, and *b* using the MATLAB function fmincon with initial values γ_0_ = 0.2, β = 1 and *b*_0_ = 0, with the constraint β ≥ 0.

### Logistic regression model

Similar to previous studies, we fitted a logistic regression of the following form to predict the choice on trial *n* based on the previous choices, previous outcomes, and interaction between previous choices and outcomes:

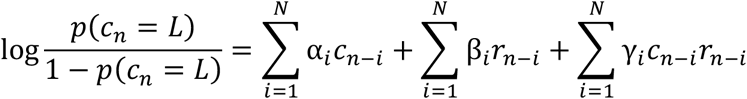

In other words, the logit was a linear combination of the previous *N* choices (*c_n-i_* = 1 for left choice and −1 for right choice), previous *N* rewards (*r_n-i_* = 1 for rewarded actions and −1 for unrewarded actions), and previous *N* interactions of choice and reward. The logistic regression model in Fig. 2g was fitted with MATLAB function mnrfit to recover the best fit coefficients α_*i*_, β_*i*_, γ_*i*_, together with the confidence intervals of these estimates. For ease of visualization, the parameters α, β and γ were normalized by their respective maximum values.

### Characterization of Q-learning and inference-based spaces

We simulated an ensemble of Q-learning and inference-based agents with parameters as described above. For each agent, the behavior was simulated for a total of 1000 blocks. To calculate the transition function of the agent, we took the average of the “signed choice”

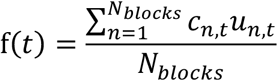

where *c_n,t_* denotes the choice in trial *t* of the block *n* (−1 for left and 1 for right choices) and *u_n,t_* denotes the unobserved hidden state in trial *t* of the block *n* (−1 for “left” state and 1 for “right” state). The signed choice ensures that *f*(*t*) is an increasing function of *t* regardless of the hidden state of the block.

The transition function *f*(*t*) was fit with a sigmoidal curve with the form

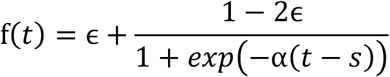

Where ϵ, α and s are free parameters of the function representing the lapse rate, slope and offset, respectively. The parameters were jointly fit with the Python function scipy.optimize.minimize(), with constraints *s* ≥ 0, *α* ≥ 0, 0≤ ϵ ≤0.5.

We also determined the foraging efficiency of the agent, *E* = *N_rewarded_/N*, where *N_rewarded_* is the number of rewarded trials and *N* is the total number of trials in the session.

### Clustering into behavioral regimes (Fig. 5)

The above fitting procedure was done for all 650 agents (25 x 20 Q-learning and 15 x 10 inference-based agents). We pooled the four behavioral features, ϵ, α, *s*, and *E*, from these agents to form a 4 x 650 feature matrix, representing 4 features/agent and 650 agents. We applied a density-based clustering method to segment the cloud of points into distinct domains. First, the four-dimensional features were embedded into a two-dimensional t-SNE space using the MATLAB tsne function with Euclidean distance metric and perplexity of 30. For the 90-10 world, the perplexity was adjusted to 25 to achieve better convergence of the t-SNE clusters.

We formed 2-D histograms of the data points in the t-SNE space using the MATLAB hist2d function (*n* = 25 bins in each dimension). These histograms were heat maps that indicated regions of high concentration of the data points. The histograms were mean-filtered by a square kernel of size 4×4, and local ‘noise’ maxima with heights less than 3 were suppressed. A watershed algorithm was run on the resulting heat map to identify the local clusters of high density. The identities of these clusters were assigned after mapping back to the location in the Q-learning or inference-based parameter spaces.

### Decoding analysis (Fig. 6)

We generated a synthetic data set using computational simulations that serve as the basis for our decoding analysis. For each agent in the Q-learning and inference-based parameter spaces, we performed repeated simulations in 50 synthetic experimental sessions with 20 block transitions per session. For each synthetic session, we obtained the transition function *f*(*t*), and fit a sigmoidal curve to estimate the four features, ϵ, α, *s*, and *E* of the behavior. The fitted slope was capped at a maximum value of 20 to avoid outliers. To balance the number of training examples for different classes in the data set, we determined the number of training examples, *n*_1_,…,*n*_6_, for each of the six classes (Q1-4, or IB5-6), and subsampled each class so that each class contains N = *min*(*n*_1_,…,*n*_6_) examples. We split this data into a training set (containing 80% of the data) and a test set (20% of the data). Each of the four features were normalized to mean 0 and standard deviation 1. A *k*-nearest neighbor (kNN) decoder was trained on the training set to predict the behavioral regime (1 to 6). Its performance was evaluated on the held-out test set. The accuracy of the decoder was measured both by the fraction of correctly labeled examples per regime, and by the Matthews Correlation Coefficient.

### Session-averaged decoding (Fig. 7)

For each behavioral session consisting of *N* blocks, we obtained the transition function *f*(*t*) as described in *Characterization of Q-learning and inference-based spaces*. We obtained the sigmoidal fit of this function and determined the parameters ϵ, α, *s*, and *E* of the session. The features were input to the kNN decoder that was trained in the *Decoding analysis* section. This results in a predicted class (Q1-4 or IB5-6) for each behavioral session. For sessions in probabilistic environments (90-10, 80-20 or 70-30), the behavioral features were input to the corresponding decoder which were trained on synthetic data from the corresponding environment type.

### BlockHMM implementation

The blockHMM inference procedure was implemented based on the existing ssm toolbox that was previously developed for a wide range of Bayesian state-space models^63^.

We added an implementation to this toolbox by specifying a new set of transition and observation probabilities which specify the blockHMM process. Each observation was defined by three vectors, ***α***, ***s*** and *ϵ* representing the parameters of the sigmoidal transition function for each of the *K* HMM modes (each vector has dimension *K* x 1). The vectors were initialized to ***α*** = 4, *s_i_* = 0.2, *ϵ_i_* = 0.3 for all 1 ≤ *i* ≤ *K*.

Given the hidden state in block *i*, i.e. given *z_i_* = *k*, the joint log likelihood of the observed choices in the block is defined via the sigmoidal transition function specified by parameters α_*k*_, *s_k_*, ϵ_*k*_

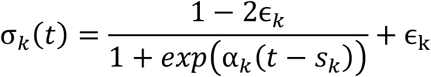

where *t* = 1, 2,…,T enumerates the position of the trials in the block.

The log-likelihood for a “signed” choice *y_t_* (the product of choice *c_t_* and hidden state *u_t_*) is that of a Bernoulli random variable with a rate of *σ_k_* (*t*).

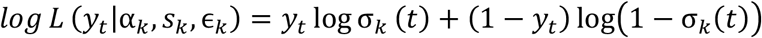

The joint log-likelihood of the observed choices in the block *i* is the sum of the log-likelihoods of individual trials

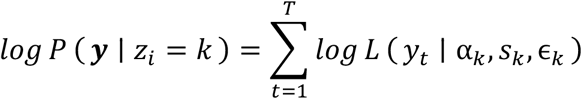

The joint log-likelihood for the whole session is the sum of the log-likelihood in individual blocks. The hidden states evolved according to a Markovian process with stationary transition governed by a transition matrix *T* with dimension *K* x *K*.

The blockHMM was fit with an Expectation-Maximization (EM) algorithm. The hidden states were initialized based on *k*-means clustering with *K* clusters. The implementation of the EM algorithm was the same as described previously for the ssm toolbox. We used the L-BFGS algorithm for the M-step when updating the values of ***α***, ***s*** and ***ϵ***, with constraints ***s*** ≥ 0.01, ***α*** ≥ 0.01,0.01 ≤ ***ϵ*** ≤ 0.5.

To evaluate the cross-validated log-likelihood (Fig. 8d), we split the data into 80% training set and 20% test set. The blockHMM was run on the training set and the log-likelihood *L_test_* was evaluated on the test set. We normalized this cross validated log-likelihood by

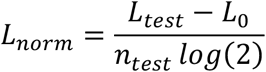

where *L_0_* is the cross-validated log-likelihood of a null model (a Bernoulli(*p*) model where *p* is the observed fraction of trials where *y_t_* = 1), *n_test_* is the number of trials in the test set.

### Synthetic agent simulation

The synthetic agent (Fig. 8c-g) was simulated with *K* = 3 HMM modes with parameters *s*_1_ = 4, *α*_1_, = 0.2, ϵ_1_ = 0.3; *s_2_* = 1, α_2_ = 0.8, ϵ_2_ = 0.15; s_3_ = 9, α_3_ = 1.5, ϵ_3_ = 0.05. The true transition matrix of the agent was

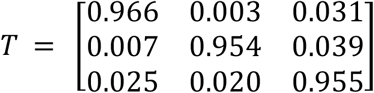

The behavior was generated for *N* = 1000 blocks, each block consisting of 30 trials.

### BlockHMM fitting to animal behavior

For each animal, we concatenated the behavioral choices from all training sessions into a *B* x *T* matrix where *B* is the total number of blocks from all the sessions and *T* = 15 is the number of trials in each block (for blocks that are longer than *T* trials we kept only the first *T* trials of that block). The blockHMM fitting procedure was run on this matrix for *K* = 1, 2, 3,…,8 modes. We chose the value of *K* that maximized the normalized log-likelihood of the test set (*L_norm_*). We capped this K value at 6 for interpretability of the model (i.e. if the value of *K* with the highest log-likelihood is higher than 6, we chose *K* = 6 as the optimal value).

After fitting the blockHMM model, we recovered parameters *s_k_*, α_*k*_, ϵ_*k*_ for individual modes in the model. We determined the foraging efficiency *E_k_* by numerically integrating the area under the curve of the choice transition function (with a step size of 0.1)

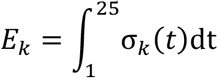

Together, the four parameters *s_k_*, α_*k*_, ϵ*_k_*, *E_k_* are input into the kNN decoder that was trained in *“Decoding analysis*” to infer the behavioral regime (Q1-4, IB5-6) of each of the HMM modes.

## Data availability

The data that support the findings of this study are available from the corresponding authors upon reasonable request.

## Code availability

Code used in this study is available at https://github.com/nhat-le/switching-simulations.

## Acknowledgements

The authors thank Tzuhsuan Ma, Morteza Sarafyazd, John Tauber, Indie Garwood and members of the Sur lab for insightful feedback on the project conceptualization and implementation of the Hidden Markov Model. This work was supported by US National Institute of Health (NIH) grants R01MH126351 and R01EY028219 (MS), K99 EB027706 (MY), Army Research Office grant W911NF-21-1-0328 (MS), Paul and Lilah Newton Brain Science Research Award (NML), and an equipment grant from the Massachusetts Life Sciences Initiative.

## Author contributions

N.M.L. conceived of the analysis framework with inputs from M.Y., M.S. and M.J. N.M.L. built the animal training apparatus. N.M.L. and M.Y. performed animal surgeries. N.M.L., M.Y., Y.W. and H.S. performed animal training. N.M.L. performed computational simulations, experimental data analyses and designed the blockHMM algorithm. M.S. and M.J. supervised the project. All authors contributed to the interpretation of the results. N.M.L. wrote the manuscript with input from all authors.

## SUPPLEMENTARY FIGURES

**Figure S1.**
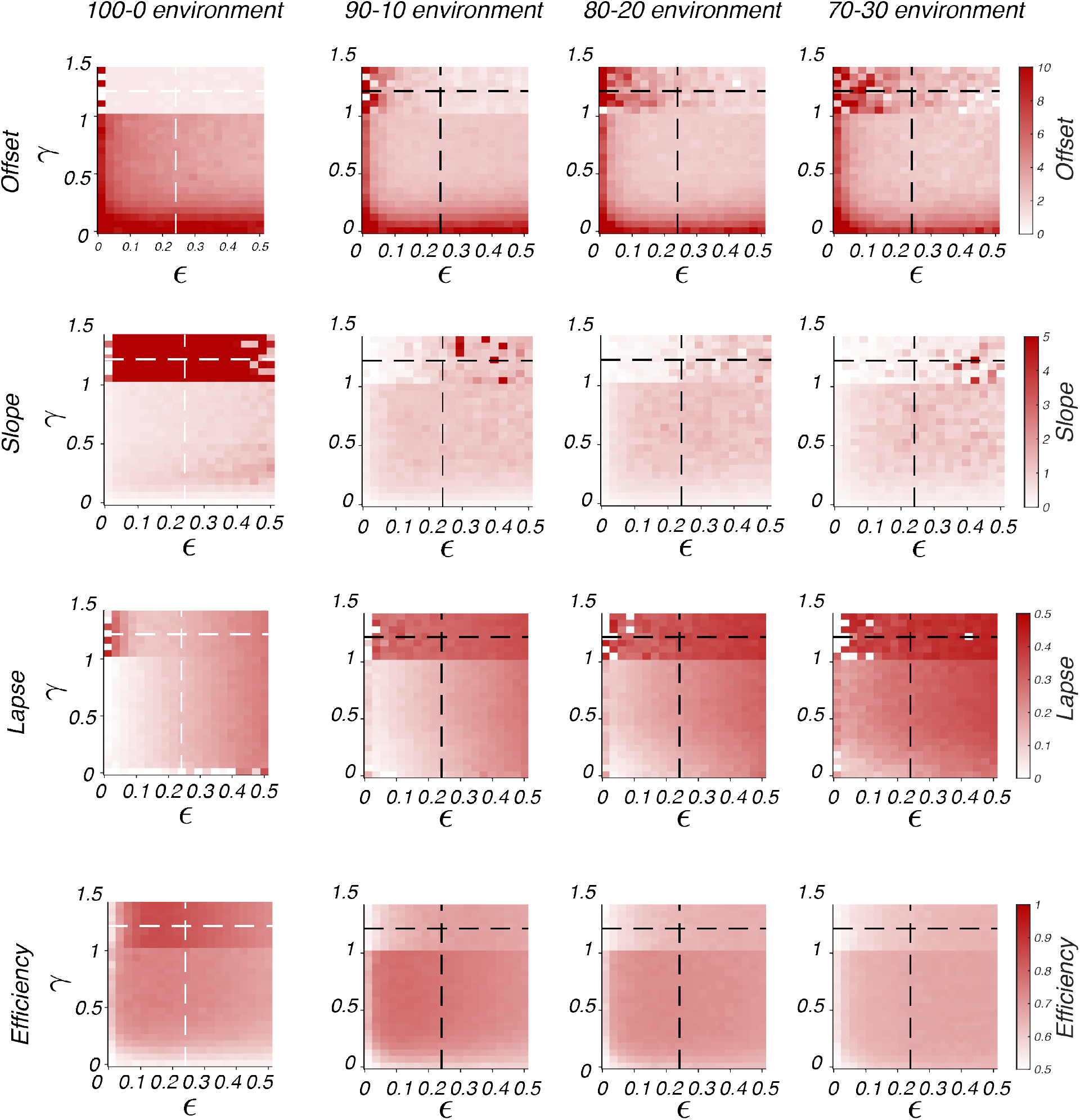
Behavioral metrics of Q-learning agents in different types of deterministic and stochastic environments (100-0, 90-10, 80-20 and 70-30). Conventions are the same as Fig. 3b.

**Figure S2.**
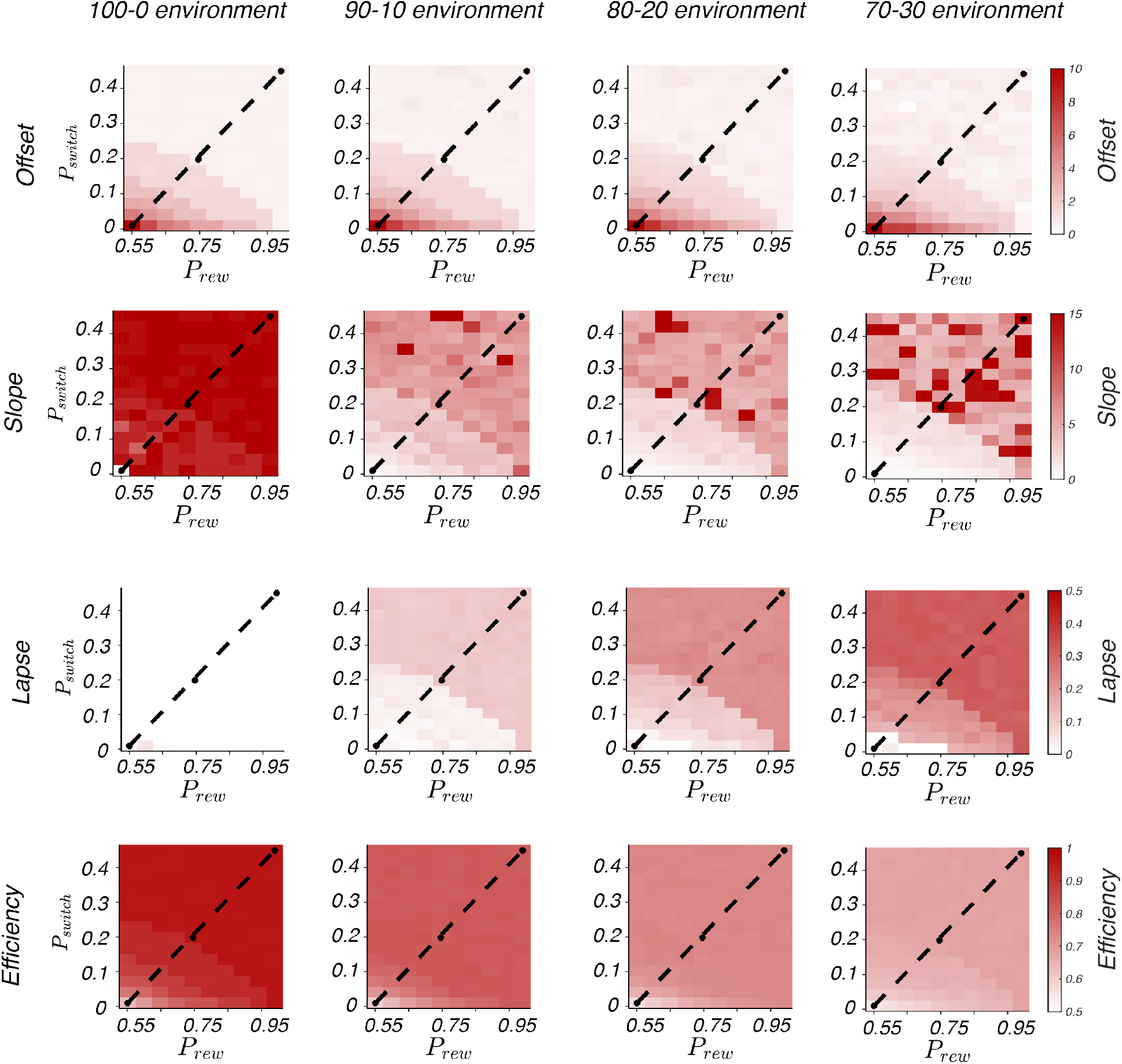
Behavioral metrics of inference-based agents in different types of deterministic and stochastic environments (100-0, 90-10, 80-20 and 70-30). Conventions are the same as Fig. 3b.

**Figure S3.**
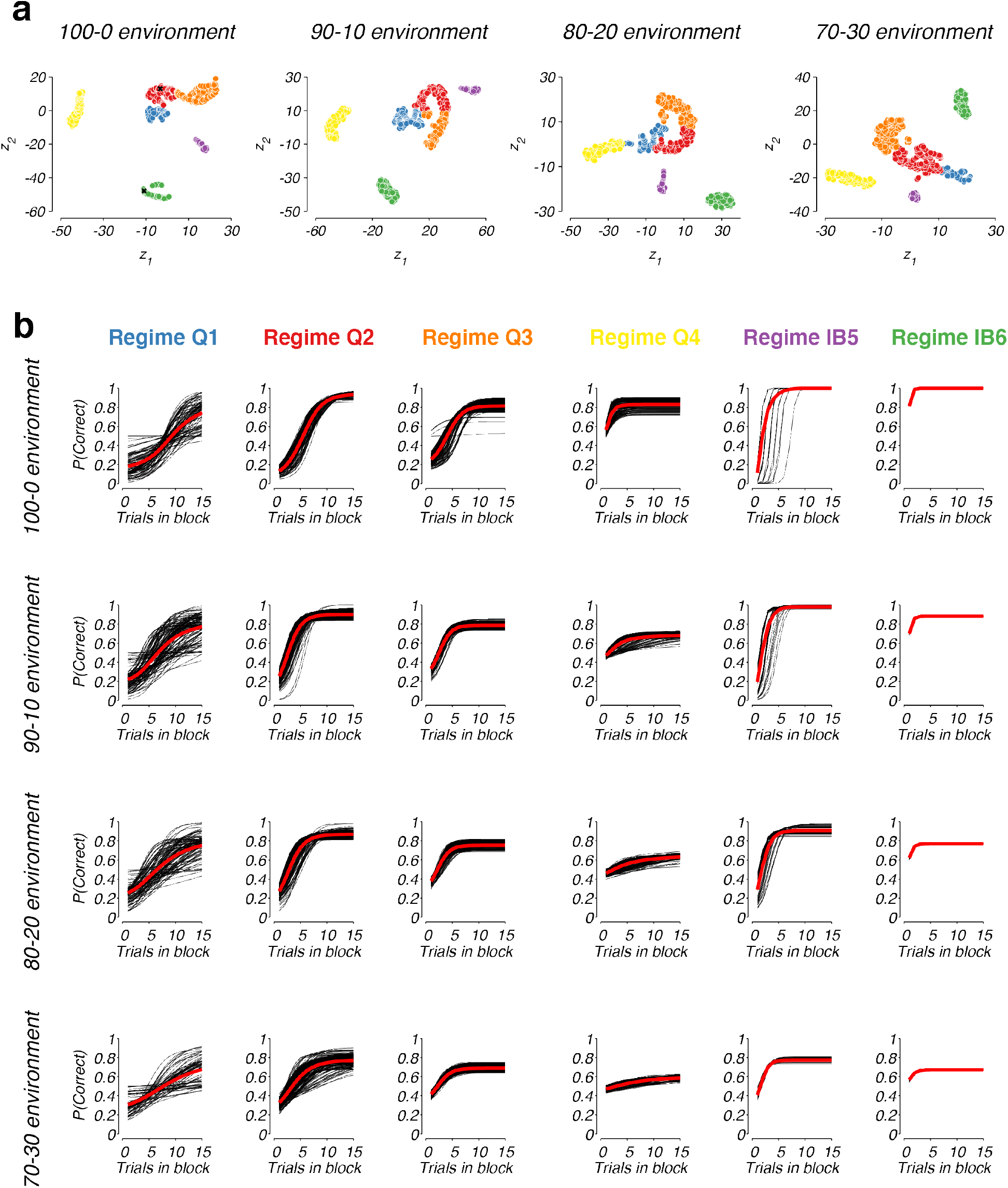
a) Non-linear embedding of all agents’ behavioral features on the t-SNE space. Points are colored based on the results of density-based segmentation (Colors of the six clusters are the same as in Fig. 5). b) Transition functions of all simulated agents grouped according to the six behavioral regimes. Red lines indicate the mean across all functions in the group.

**Figure S4.**
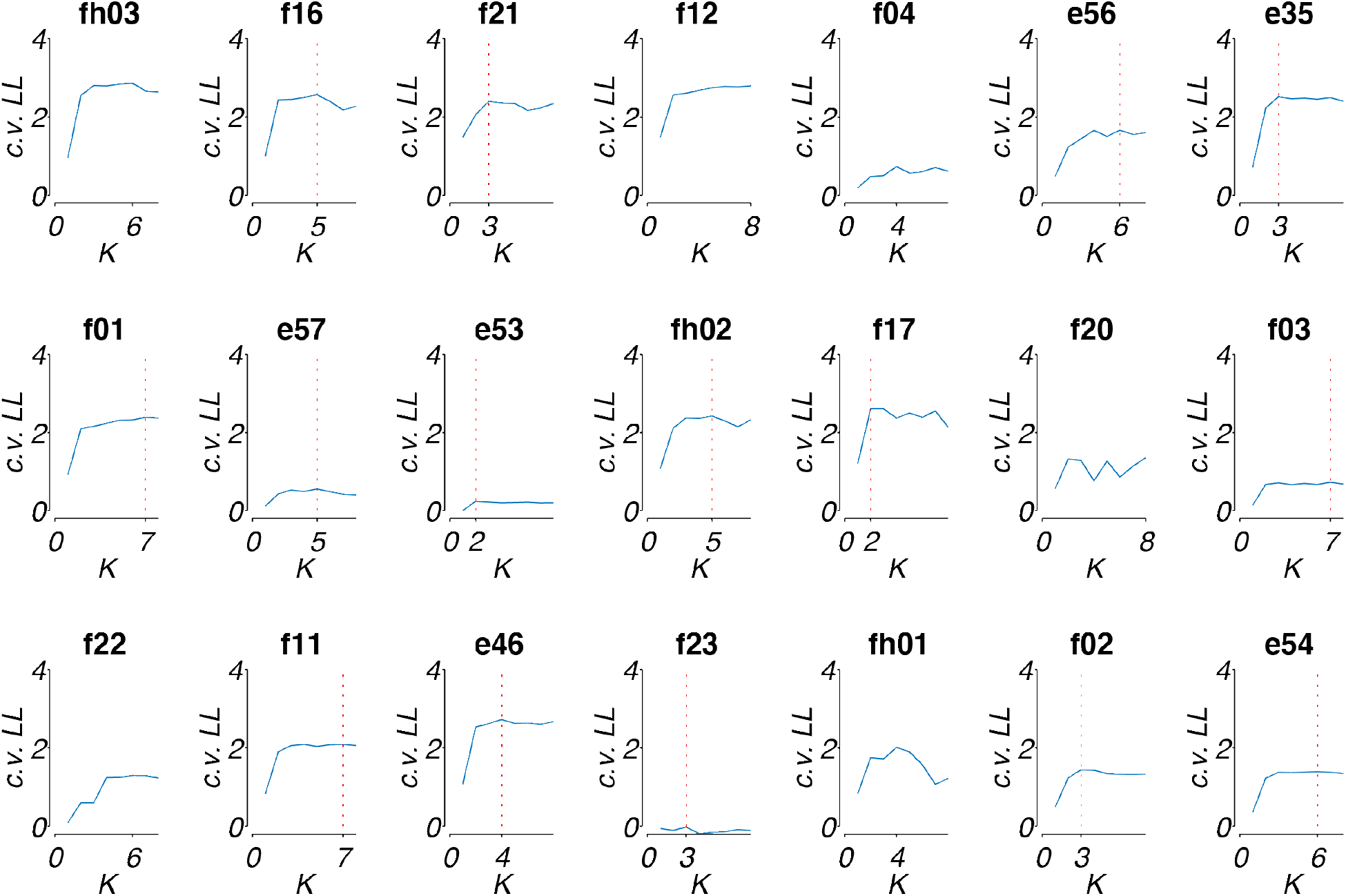
Normalized cross-validated log-likelihood for different values of *K*, the number of clusters of the blockHMM for the *n* = 21 mice used in the paper. For each animal, the value of *K* that gave the highest cross-validated log-likelihood was chosen for subsequent analyses and fitting (this *K* value is indicated by the vertical red line).

**Figure S5.**
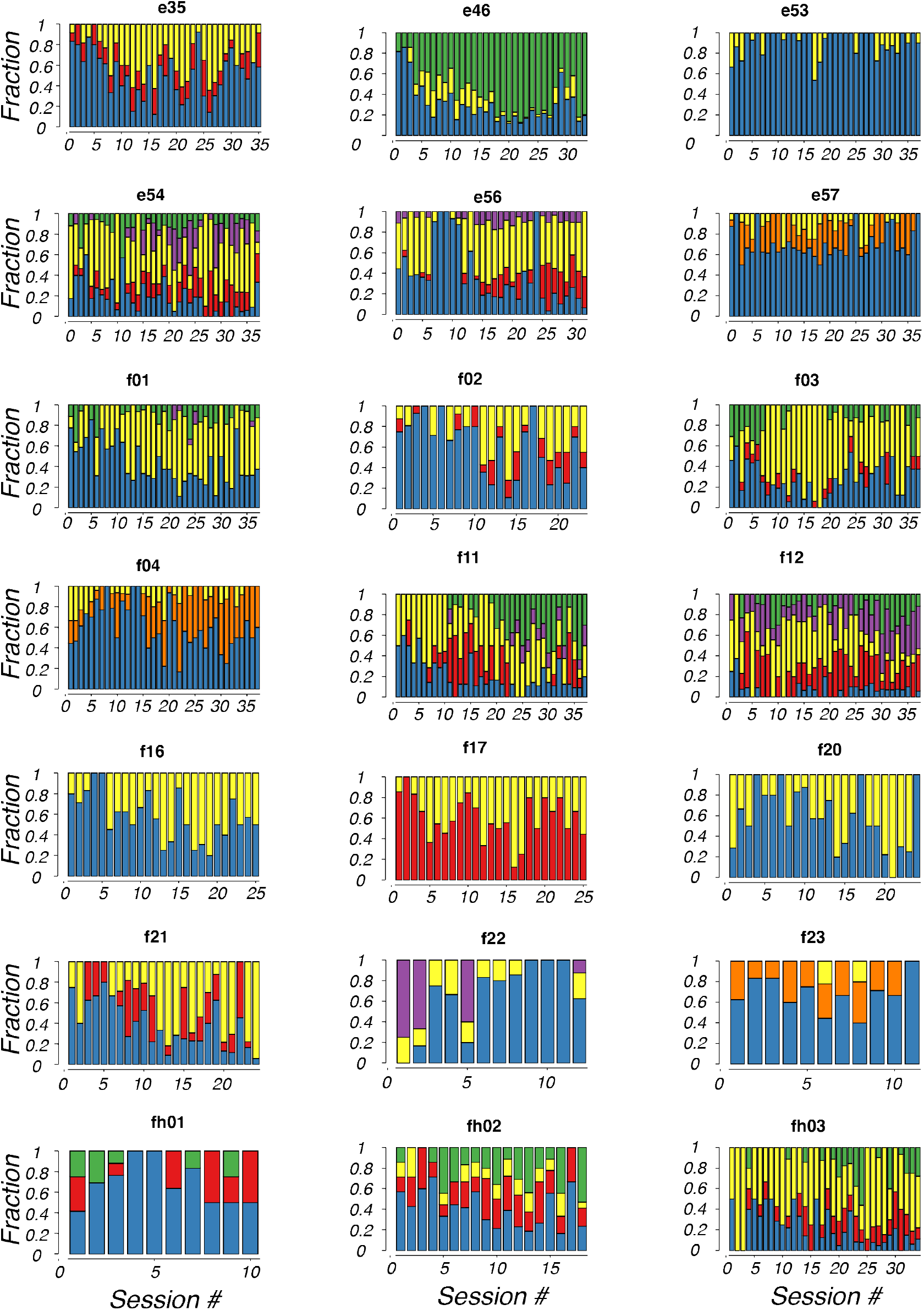
Evolution of mixture of behavioral strategies as inferred by blockHMM for all the *n* = 21 mice through different training sessions. Colors and conventions are the same as Fig. 9.

**Fig. S6.**
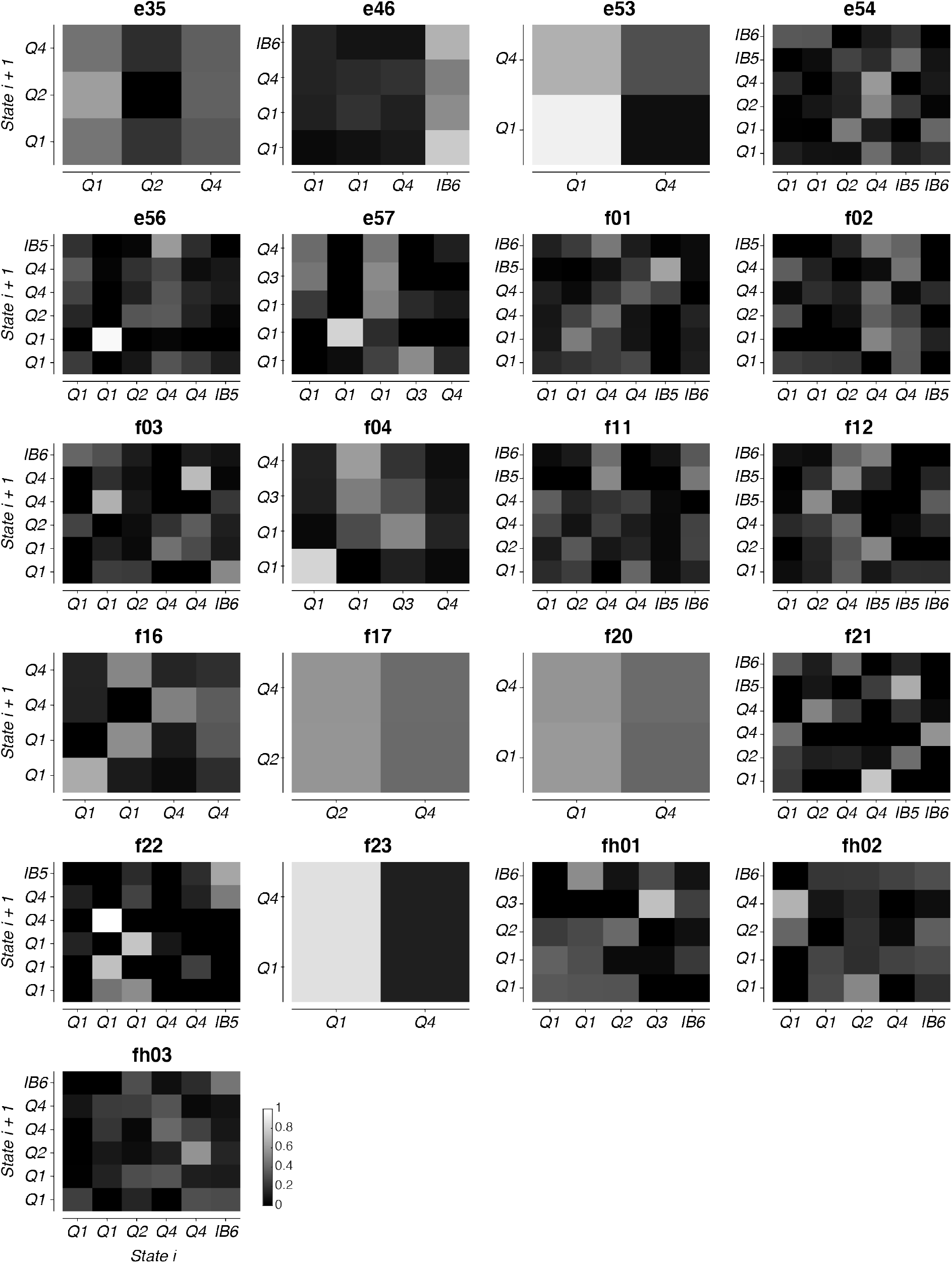
Transition functions as fitted by the blockHMM procedure for all the *n* = 21 mice analyzed in the paper.

**Fig. S7.**
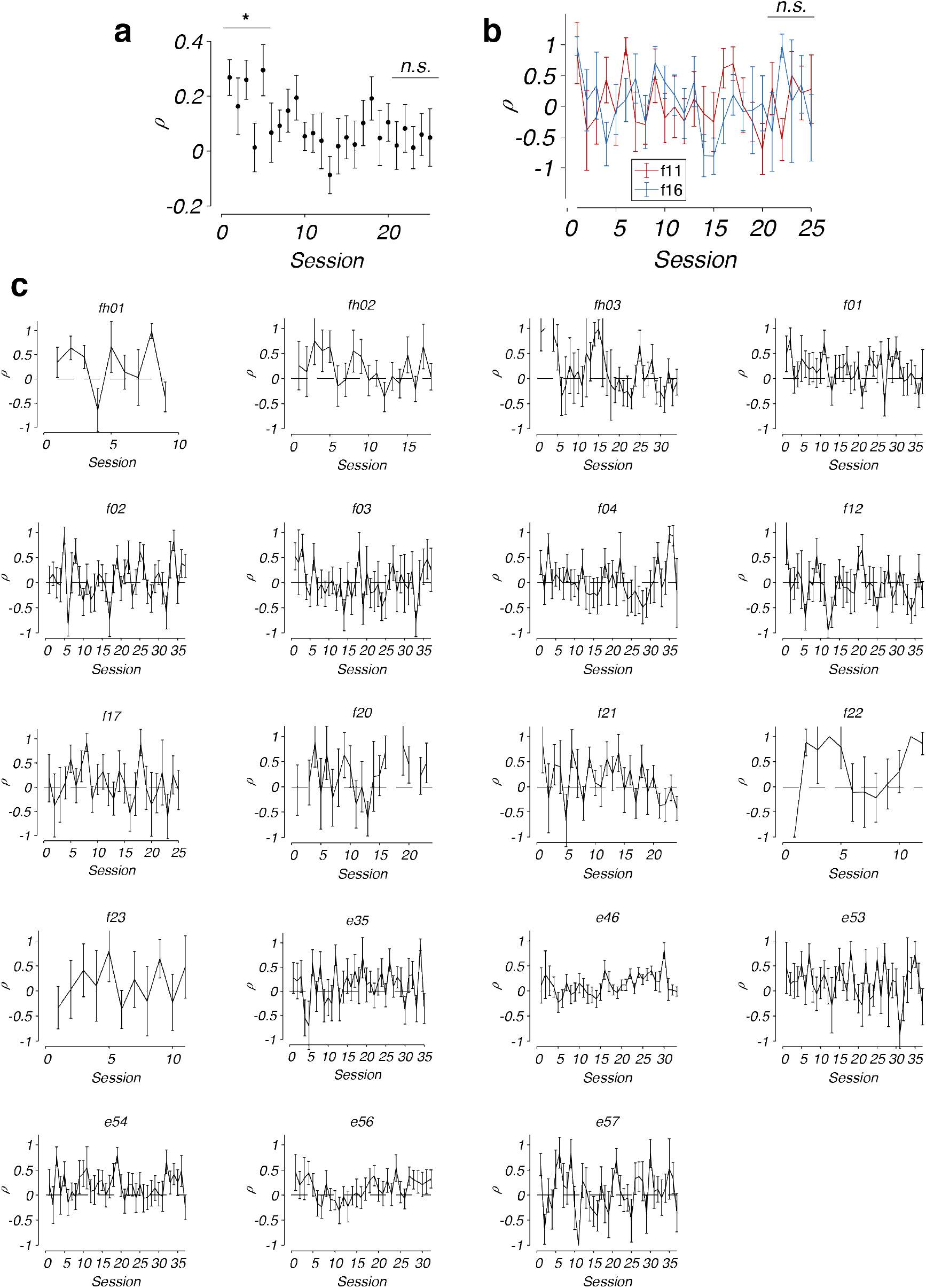
a) Average evolution of *ρ* across all experimental animals (mean ± standard errors, *n* = 21 animals). b) Comparison of the evolution of *ρ* for two animals, f11 and f16 (mean ± standard errors). c) Fitting of *ρ* for the remaining 19 animals over the course of training (mean ± standard errors).

## REFERENCES

1. Sutton, R. S. & Barto, A. G. Reinforcement learning: An introduction. (MIT press, 2018).

2. Kaelbling, L. P., Littman, M. L. & Cassandra, A. R. Planning and acting in partially observable stochastic domains. Artif. Intell. 101, 99–134 (1998).

3. Rescorla, R. A. A theory of Pavlovian conditioning: Variations in the effectiveness of reinforcement and nonreinforcement. Curr. Res. theory 64–99 (1972).

4. Watkins, C. J. C. H. & Dayan, P. Q-learning. Mach. Learn. 8, 279–292 (1992).

5. Herrnstein, R. J. Secondary reinforcement and rate of primary reinforcement. J. Exp. Anal. Behav. 7, 27–36 (1964).

6. Thompson, W. R. On the likelihood that one unknown probability exceeds another in view of the evidence of two samples. Biometrika 25, 285–294 (1933).

7. Samejima, K., Ueda, Y., Doya, K. & Kimura, M. Neuroscience: Representation of action-specific reward values in the striatum. Science (80-.). 310, 1337–1340 (2005).

8. Sugrue, L. P., Corrado, G. S. & Newsome, W. T. Matching behavior and the representation of value in the parietal cortex. Science (80-.). 304, 1782–1787 (2004).

9. Pisupati, S., Chartarifsky-Lynn, L., Khanal, A. & Churchland, A. K. Lapses in perceptual decisions reflect exploration. Elife 10, e55490 (2021).

10. Gershman, S. J. Deconstructing the human algorithms for exploration. Cognition 173, 34–42 (2018).

11. Wilson, R. C., Geana, A., White, J. M., Ludvig, E. A. & Cohen, J. D. Humans use directed and random exploration to solve the explore–exploit dilemma. J. Exp. Psychol. Gen. 143, 2074 (2014).

12. Trepka, E. et al. Entropy-based metrics for predicting choice behavior based on local response to reward. Nat. Commun. 12, 6567 (2021).

13. Schultz, W. Predictive reward signal of dopamine neurons. J. Neurophysiol. 80, 1–27 (1998).

14. Lau, B. & Glimcher, P. W. Value Representations in the Primate Striatum during Matching Behavior. Neuron 58, 451–463 (2008).

15. Hattori, R., Danskin, B., Babic, Z., Mlynaryk, N. & Komiyama, T. Area-Specificity and Plasticity of History-Dependent Value Coding During Learning. Cell 177, 1858–1872.e15 (2019).

16. Hattori, R. & Komiyama, T. Context-dependent persistency as a coding mechanism for robust and widely distributed value coding. Neuron 110, 502–515.e11 (2022).

17. Vertechi, P. et al. Inference-Based Decisions in a Hidden State Foraging Task: Differential Contributions of Prefrontal Cortical Areas. Neuron 106, 166–176.e6 (2020).

18. Costa, V. D., Tran, V. L., Turchi, J. & Averbeck, B. B. Reversal learning and dopamine: a bayesian perspective. J. Neurosci. 35, 2407–2416 (2015).

19. Beron, C. C., Neufeld, S. Q., Linderman, S. W. & Sabatini, B. L. Efficient and stochastic mouse action switching during probabilistic decision making. bioRxiv (2021).

20. Cox, J. & Witten, I. B. Striatal circuits for reward learning and decision-making. Nat. Rev. Neurosci. 20, 482–494 (2019).

21. Yin, H. H., Knowlton, B. J. & Balleine, B. W. Inactivation of dorsolateral striatum enhances sensitivity to changes in the action–outcome contingency in instrumental conditioning. Behav. Brain Res. 166, 189–196 (2006).

22. Tai, L. H., Lee, A. M., Benavidez, N., Bonci, A. & Wilbrecht, L. Transient stimulation of distinct subpopulations of striatal neurons mimics changes in action value. Nat. Neurosci. 15, 1281–1289 (2012).

23. Verharen, J. P. H., Adan, R. A. H. & Vanderschuren, L. J. M. J. Differential contributions of striatal dopamine D1 and D2 receptors to component processes of value-based decision making. Neuropsychopharmacology 44, 2195–2204 (2019).

24. Donahue, C. H., Liu, M. & Kreitzer, A. C. Distinct value encoding in striatal direct and indirect pathways during adaptive learning. bioRxiv 277855 (2018).

25. Angela, J. Y. & Dayan, P. Uncertainty, neuromodulation, and attention. Neuron 46, 681–692 (2005).

26. Sarafyazd, M. & Jazayeri, M. Hierarchical reasoning by neural circuits in the frontal cortex. Science (80-.). 364, (2019).

27. Miller, K. J., Botvinick, M. M. & Brody, C. D. From predictive models to cognitive models: Separable behavioral processes underlying reward learning in the rat. bioRxiv 461129 (2021).

28. Roy, N. A., Bak, J. H., Akrami, A., Brody, C. D. & Pillow, J. W. Extracting the dynamics of behavior in sensory decision-making experiments. Neuron 109, 597–610.e6 (2021).

29. Ashwood, Z. C. et al. Mice alternate between discrete strategies during perceptual decision-making. Nat. Neurosci. 1–12 (2022).

30. Cazettes, F., Murakami, M., Renart, A. & Mainen, Z. F. Reservoir of decision strategies in the mouse brain. (2021).

31. Calhoun, A. J. & Hayden, B. Y. The foraging brain. Curr. Opin. Behav. Sci. 5, 24–31 (2015).

32. Witten, I. H. The apparent conflict between estimation and control—A survey of the two-armed bandit problem. J. Franklin Inst. 301, 161–189 (1976).

33. Ma, W. J. Bayesian Decision Models: A Primer. Neuron 104, 164–175 (2019).

34. Steinmetz, N. A., Zatka-Haas, P., Carandini, M. & Harris, K. D. Distributed coding of choice, action and engagement across the mouse brain. Nature 576, 266–273 (2019).

35. Kheifets, A. & Gallistel, C. R. Mice take calculated risks. Proc. Natl. Acad. Sci. 109, 8776–8779 (2012).

36. Lak, A. et al. Reinforcement biases subsequent perceptual decisions when confidence is low, a widespread behavioral phenomenon. Elife 9, e49834 (2020).

37. Ito, M. & Doya, K. Validation of decision-making models and analysis of decision variables in the rat basal ganglia. J. Neurosci. 29, 9861–9874 (2009).

38. Berman, G. J., Bialek, W. & Shaevitz, J. W. Predictability and hierarchy in Drosophila behavior. Proc. Natl. Acad. Sci. 113, 11943–11948 (2016).

39. Linderman, S., Nichols, A., Blei, D., Zimmer, M. & Paninski, L. Hierarchical recurrent state space models reveal discrete and continuous dynamics of neural activity in C. elegans. BioRxiv 621540 (2019).

40. Buchanan, E. K., Lipschitz, A., Linderman, S. W. & Paninski, L. Quantifying the behavioral dynamics of C. elegans with autoregressive hidden Markov models. in Workshop on Worm’s neural information processing at the 31st conference on neural information processing systems (2017).

41. Grossman, C. D., Bari, B. A. & Cohen, J. Y. Serotonin neurons modulate learning rate through uncertainty. Curr. Biol. 32, 586–599.e7 (2022).

42. Bari, B. A. et al. Stable Representations of Decision Variables for Flexible Behavior. Neuron 103, 922–933.e7 (2019).

43. Bloem, B., Huda, R., Sur, M. & Graybiel, A. M. Two-photon imaging in mice shows striosomes and matrix have overlapping but differential reinforcement-related responses. Elife 6, e32353 (2017).

44. Eckstein, M. K., Master, S. L., Dahl, R. E., Wilbrecht, L. & Collins, A. G. E. The Unique Advantage of Adolescents in Probabilistic Reversal: Reinforcement Learning and Bayesian Inference Provide Adequate and Complementary Models. BioRxiv 2007–2020 (2021).

45. Daw, N. D., Gershman, S. J., Seymour, B., Dayan, P. & Dolan, R. J. Model-based influences on humans’ choices and striatal prediction errors. Neuron 69, 1204–1215 (2011).

46. Akam, T. et al. The Anterior Cingulate Cortex Predicts Future States to Mediate Model-Based Action Selection. Neuron 109, 149–163.e7 (2021).

47. Lee, S. W., Shimojo, S. & O’Doherty, J. P. Neural Computations Underlying Arbitration between Model-Based and Model-free Learning. Neuron 81, 687–699 (2014).

48. Haith, A. M. & Krakauer, J. W. Model-Based and Model-Free Mechanisms of Human Motor Learning BT - Progress in Motor Control. in (eds. Richardson, M. J., Riley, M. A. & Shockley, K.) 1–21 (Springer New York, 2013).

49. Niv, Y. Learning task-state representations. Nat. Neurosci. 22, 1544–1553 (2019).

50. Daw, N. D. Trial-by-trial data analysis using computational models. Decis. making, Affect. Learn. Atten. Perform. XXIII 23, (2011).

51. Rosenberg, M., Zhang, T., Perona, P. & Meister, M. Mice in a labyrinth show rapid learning, sudden insight, and efficient exploration. Elife 10, e66175 (2021).

52. Fonio, E., Benjamini, Y. & Golani, I. Freedom of movement and the stability of its unfolding in free exploration of mice. Proc. Natl. Acad. Sci. 106, 21335–21340 (2009).

53. Gordon, G., Fonio, E. & Ahissar, E. Emergent exploration via novelty management. J. Neurosci. 34, 12646–12661 (2014).

54. Thompson, S. M., Berkowitz, L. E. & Clark, B. J. Behavioral and neural subsystems of rodent exploration. Learn. Motiv. 61, 3–15 (2018).

55. Wichmann, F. A. & Hill, N. J. The psychometric function: I. Fitting, sampling, and goodness of fit. Percept. Psychophys. 63, 1293–1313 (2001).

56. Erlich, J. C., Brunton, B. W., Duan, C. A., Hanks, T. D. & Brody, C. D. Distinct effects of prefrontal and parietal cortex inactivations on an accumulation of evidence task in the rat. Elife 4, e05457 (2015).

57. Carandini, M. & Churchland, A. K. Probing perceptual decisions in rodents. Nat. Neurosci. 16, 824–831 (2013).

58. Tijsma, A. D., Drugan, M. M. & Wiering, M. A. Comparing exploration strategies for Q-learning in random stochastic mazes. in 2016 IEEE Symposium Series on Computational Intelligence (SSCI) 1–8 (IEEE, 2016).

59. Thrun, S. B. Efficient exploration in reinforcement learning. (1992).

60. Collins, A. G. E. & Frank, M. J. How much of reinforcement learning is working memory, not reinforcement learning? A behavioral, computational, and neurogenetic analysis. Eur. J. Neurosci. 35, 1024–1035 (2012).

61. Bhagat, J., Wells, M. J., Harris, K. D., Carandini, M. & Burgess, C. P. Rigbox: an Open-Source toolbox for probing neurons and behavior. Eneuro 7, (2020).

62. Burgess, C. P. et al. High-Yield Methods for Accurate Two-Alternative Visual Psychophysics in Head-Fixed Mice. Cell Rep. 20, 2513–2524 (2017).

63. Linderman, S., Antin, B., Zoltowski, D. & Glaser, J. SSM: Bayesian Learning and Inference for State Space Models. (2020).

